# Genetics of *trans*-regulatory variation in gene expression

**DOI:** 10.1101/208447

**Authors:** Frank W. Albert, Joshua S. Bloom, Jake Siegel, Laura Day, Leonid Kruglyak

## Abstract

Heritable variation in gene expression provides a critical bridge between differences in genome sequence and the biology of many traits, including common human diseases. However, the sources of most regulatory genetic variation remain unknown. Here, we used transcriptome profiling in 1,012 yeast segregants to map the genetic basis of variation in gene expression with high statistical power. We identified expression quantitative trait loci (eQTL) that together account for over 70% of the total genetic contribution to variation in mRNA levels, allowing us to examine the sources of regulatory variation comprehensively. We found that variation in the expression of a typical gene has a complex genetic architecture involving multiple eQTL. We also detected hundreds of eQTL pairs with significant non-additive interactions in an unbiased genome-wide scan. Although most genes were influenced by a local eQTL located close to the gene, most expression variation arose from distant, *trans*-acting eQTL located far from their target genes. Nearly all distant eQTL clustered at 102 “hotspot” locations, some of which influenced the expression of thousands of genes. Hotspot regions were enriched for transcription factor genes and altered expression of their target genes though both direct and indirect mechanisms. Many local eQTL had no detectable effects on the expression of other genes in *trans*. These results reveal the complexity of genetic influences on transcriptome variation in unprecedented depth and detail.

## Main text

Differences in gene expression among individuals arise in part from DNA sequence differences in regulatory elements and in regulatory genes. Regions of the genome that contain regulatory variants can be identified by tests of genetic linkage or association between mRNA levels and DNA polymorphisms in large collections of individuals. Regions for which such tests show statistical significance are known as eQTL^1^. Regulatory variation is widespread in the species for which it has been studied; indeed, in humans, the expression of nearly every gene appears to be influenced by one or more eQTL^2,3^.

In humans, eQTL are typically mapped by genome-wide association studies (GWAS) in unrelated individuals. To cover the genome, human GWAS must test a very large number of variants, resulting in a high multiple-testing burden and low statistical power. As a result, most human eQTL GWAS have been limited to searches for “local” eQTL that are located close to the genes they influence^4,5^. The power to detect local eQTL is higher because focused local tests reduce the multiple-testing burden, and because local eQTL tend to have larger effect sizes. However, genome-wide estimates show that most regulatory variation does not arise from local eQTL. Instead, it arises from “distant” eQTL, which are located far from the genes they influence, typically on different chromosomes, and which exert their effects through trans-acting factors^6,7^. Although a limited number of *trans*-acting human eQTL has been discovered^2,3,6–13^, the vast majority remains unknown. As a consequence, we know relatively little about this crucial source of regulatory genetic variation.

In model organisms, eQTL can be identified by linkage analysis in panels of offspring obtained from crosses of genetically different individuals^14^. In this design, longer blocks of linkage reduce the number of statistical tests required to cover the genome. As a result, many local and distant eQTL have been discovered in such studies. However, even in model organisms, sample size limitations have to date resulted in insufficient statistical power to detect most eQTL. This limitation has manifested itself as “missing heritability”: detected eQTL tend to account for only a fraction of the measured heritable component of gene expression variation. Here, we address this limitation by carrying out an eQTL study in a very large panel of segregants from a cross between two yeast strains. The high power of our study allowed us to identify eQTL that account for the great majority of heritable expression variation in this cross, and to characterize the distant component of regulatory variation in unprecedented detail.

### Deep eQTL mapping explains most gene expression heritability

We developed an experimental pipeline for high-throughput generation of RNA-seq data in yeast and obtained high-quality expression measurements (Supplementary Data 1 & 2) for 5,720 genes in 1,012 segregants from a cross between laboratory and wine yeast strains (hereafter, BY and RM, respectively). We obtained high-confidence genotypes at 11,530 variant sites from low-coverage whole-genome sequences of the segregants^15^ (Supplementary Data 3). We used the genotype and RNA-seq data for eQTL mapping and identified 36,498 eQTL for 5,643 genes at a false discovery rate (FDR) of 5% (Supplementary Data 4). Only 77 genes had no detected eQTL. Among the genes with at least one detected eQTL, the median number was 6, with a maximum of 21 (Figure 1A, Supplementary Discussion 1). Previous eQTL mapping in 112 segregants from this cross detected an average of less than one eQTL per gene as a consequence of much lower statistical power^14,16^. That data set was used to obtain indirect estimates of the distribution of the number of eQTL per gene^17^, and these agree closely with the distribution of directly detected eQTL observed in the current study. The observed distribution of the number of loci also closely matched the distribution we reported for loci influencing 160 protein levels studied with the highly powered X-pQTL approach^18^. Our results provide direct demonstration that variation in expression levels of nearly all genes has a complex genetic basis.

**Figure 1.**
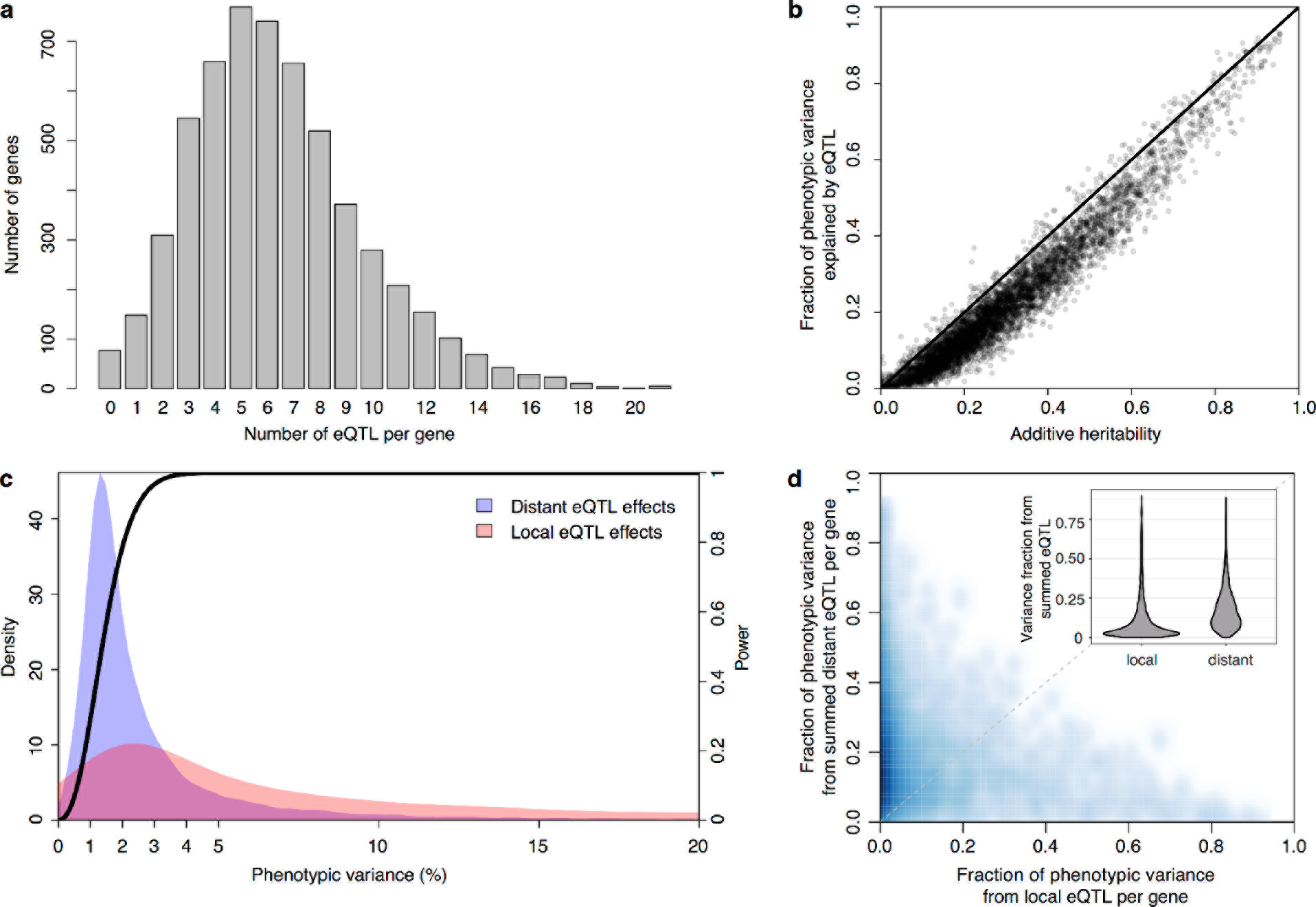
Detection and effects of local and distant eQTL. a) Histogram showing the number of eQTL per gene. b) Most additive heritability for transcript abundance variation is explained by detected eQTL. The total variance explained by detected eQTL for each transcript (y-axis) is plotted against the additive heritability (h^2^). The diagonal line represents a scenario under which the variance explained by eQTL exactly matches the heritability; the line is shown as a visual guide. c) Power to detect eQTL as a function of effect size, and distributions of observed local and distant eQTL effects. The black curve corresponds to the statistical power (right y-axis) for eQTL detection at a genome-wide significance threshold. Colored areas show the density of individual significant eQTL (left y-axis) that explain a given fraction of phenotypic variance (x-axis) for distant (blue) and local (red) eQTL. Note that the x-axis is truncated at 20% variance explained to aid visualization of smaller effects, and omits a long tail of rare, large eQTL. d) Scatterplot showing the fraction of phenotypic variance explained by the local eQTL (x-axis) and the fraction of variance explained by the sum of the distant eQTL for each gene (y-axis). The diagonal line represents the case of equal local and distant variance fractions, and is shown as a visual guide. Darker shades of blue indicate a higher density of genes. Inset: Violin plots of the distributions of fractions of phenotypic variance explained by summed local and distant eQTL, respectively.

We used our data to estimate the additive heritability of the expression level of each gene (i.e., the fraction of expression variance attributable to genetic factors; Supplementary Figure 1; Supplementary Data 5). We observed a median heritability of 26%, with a maximum of 95%. Our estimates are consistent with studies of gene expression in humans^6,7,19,20^, but are lower than those typically seen for organismal traits in this cross^15,21^, suggesting a greater contribution of environmental and stochastic factors to gene expression variation. Across genes, heritability was positively correlated with mean expression and with expression variance, and negatively correlated with the number of protein-protein and synthetic genetic interaction partners, as well as with gene essentiality (p ≤ 0.005) (Supplementary Figure 2; Supplementary Discussion 2 & 3; Supplementary Table 1; Supplementary Data 6).

In contrast to previous eQTL studies, the detected eQTL explained most of the estimated additive gene expression heritability (a median across genes of 71.5%) (Figure 1B, 10-fold crossvalidation). Low missing heritability in our data is explained by the high power of our experiment. We had greater than 90% power to detect eQTL that explain at least 2.5% of expression variance (Figure 1C). The distribution of effect sizes of detected eQTL is strongly weighted toward small effects (median 1.9% of variance explained; Figure 1C), suggesting that the remaining missing heritability is explained by undetected eQTL with even smaller effects. These results are consistent with those observed for organismal traits in this cross^15,21^. Thus, we have discovered most eQTL with substantial effects that segregate in this cross, and these jointly account for the great majority of the observed genetic variation in the transcriptome.

### Genetic expression variation arises primarily from trans-acting hotspots

We found that 2,884 genes (50% of 5,720 expressed genes) had a local eQTL (defined as an eQTL whose confidence interval includes the gene it influences) at genome-wide significance. This number rose to 4,241 genes (74% of expressed genes) when we performed eQTL analysis with only one nearby marker per gene in order to reduce the multiple testing burden (FDR < 5%). Thus, the single pair of yeast isolates used here harbors sufficient local regulatory variation to alter the expression of more than half the genes in the genome. Comparisons with allele-specific expression data^22^ support previous results^23,24^ that most but not all local eQTL act in *cis* (Supplementary Figures 3 & 4; Supplementary Discussion 4; Supplementary Tables 2 – 4; Supplementary Data 7).

The vast majority of the genome-wide significant eQTL did not overlap the genes they influenced (92%; 33,529 of 36,498); indeed, 86% were located on a different chromosome. Nearly every expressed gene (98%; 5,606) had at least one such distant, *trans*-acting eQTL. The individual effect sizes of the *trans* eQTL were smaller than those of local eQTL (median variance explained 2.8-fold less, T-test p < 2.2e-16; Figure 1C). However, for the 2,846 genes that had both a local eQTL and at least one distant eQTL, the aggregate effect of the distant eQTL per gene was larger than that of the local eQTL (median 2.6-fold more variance explained; paired T-test p < 2.2e-16; Figure 1D). Genome-wide estimates in humans have similarly indicated 1.8-fold^7^ to 3.4-fold^6^ more genetic variation arising in *trans* than in *cis*. Our results directly demonstrate the importance of *trans* acting variation by showing that mapped *trans* eQTL influence the expression of a larger number of genes and jointly contribute more to expression variance than local eQTL.

The *trans* eQTL were not uniformly distributed across the genome (Figure 2A). Instead, they clustered at 102 hotspot loci, each of which affected the expression of many genes^14,25,26^ (Figure 2B). These hotspots contained over 90% of all *trans* eQTL. The eQTL that mapped outside of the hotspots also clustered more than expected by chance (randomization p < 0.001), suggesting the existence of additional hotspots that affect the expression of too few genes to pass the stringent criteria used to define the set of 102. Isolated *trans*-acting loci that affect the expression of one or a few genes appear to be uncommon.

**Figure 2.**
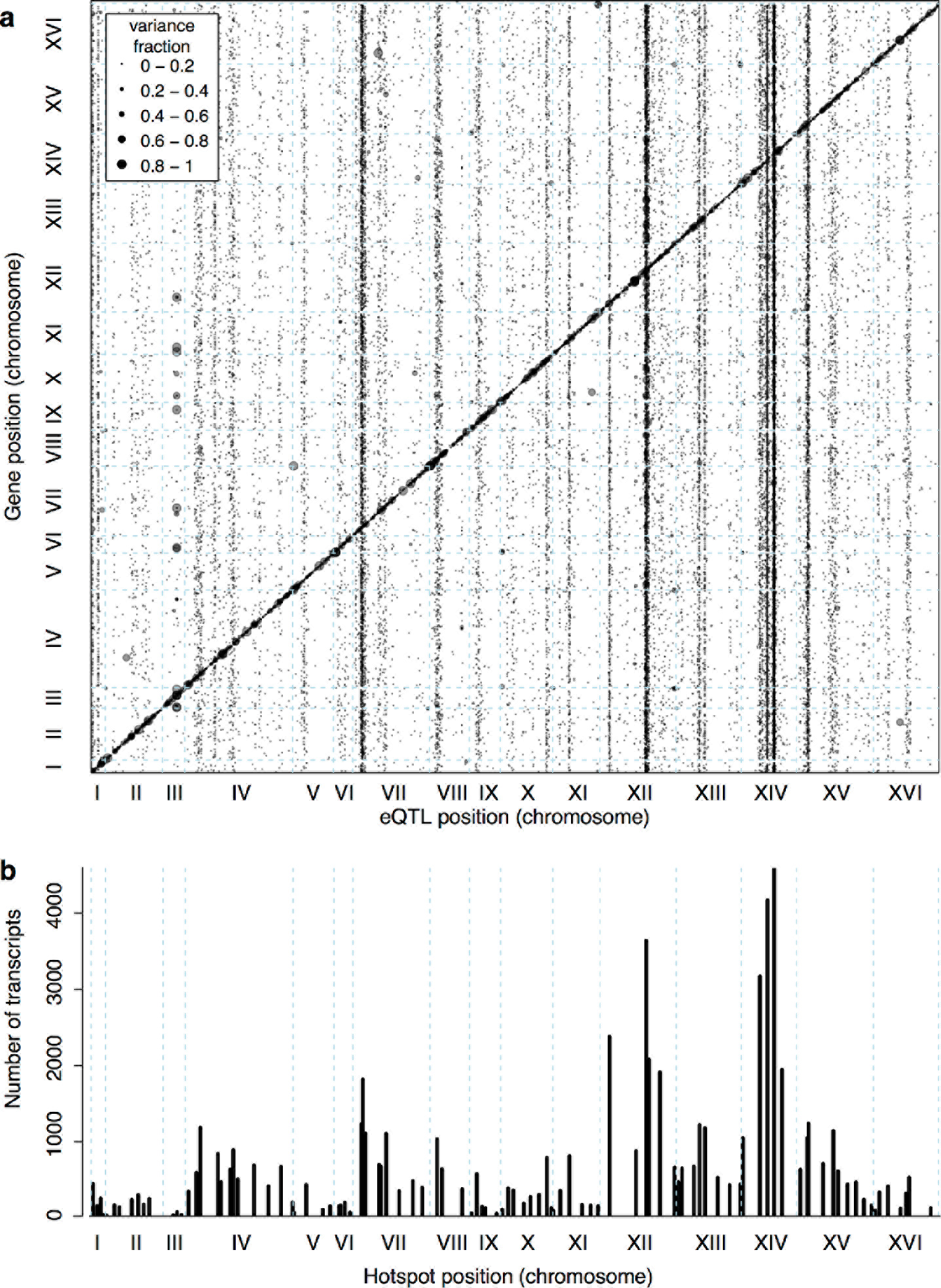
Locations of eQTL in the genome. a) Map of local and distant eQTL. The genomic locations of eQTL peaks (x-axis) are plotted against the genomic locations of the genes whose expression they influence (y-axis). The strong diagonal band corresponds to local eQTL. The many vertical bands correspond to eQTL hotspots. Point size is scaled as a function of eQTL effect size, measured in fraction of phenotypic variance explained. b) The number of gene expression traits linking to each of 102 identified eQTL hotspots (Methods) are shown as vertical bars.

The 102 hotspots affected a median of 425 genes, ranging from 26 (a newly discovered hotspot at position 166,390 bp on chromosome III) to 4,594 at the previously reported *MKT1* hotspot^27^(82% of 5,629 genes with any signal at a hotspot; Figure 2B). Three additional hotspots each affected more than half of all genes. They include a previously described hotspot at the *HAP1* gene^14^ (3,640 genes affected), as well as two newly detected hotspots on chromosome XIV. A hotspot at 372,376 bp affected 4,172 genes and is likely caused by a variant that recently arose in the *KRE33* gene in the RM parent used in our cross^28^. A hotspot at position 267,161 bp affected 3,169 genes and spans the genes *GCR2*, YNL198C, *WHI3* and *SLZ1*. These results indicate that hotspots can have extraordinarily wide-reaching effects on the transcriptome, with some influencing the expression of the majority of all genes.

Such widespread effects caused by single loci likely arise from a cascade of effects in which strong primary effects spread through the cellular regulatory network. For example, the BY allele of the transcriptional activator *HAP1* carries a transposon insertion that reduces *HAP1* function. As expected, the BY allele strongly reduced the expression of known transcriptional targets of *HAP1:* 26 out of the 69 *HAP1* targets present in our data were among the 50 genes with the largest reduction in expression in segregants carrying the BY allele of *HAP1* (p < 2.2e-16, odds ratio = 138). In total, only 75 direct transcriptional *HAP1* targets are known. Unless previous work missed thousands of *HAP1* targets, the vast majority of the 3,640 *trans* eQTL at *HAP1* must reflect indirect, secondary consequences of the direct transcriptional effects. *HAP1* is an activator of genes involved in cellular respiration. Thus, the many secondary effects of the BY *HAP1* allele on gene expression may be mediated by cellular responses to altered metabolism arising from reduced respiration.

### Causal genes underlying hotspots

Functional analysis of eQTL hotspots requires identification of the underlying causal genes, and this has been challenging to do systematically. We developed a multivariate fine-mapping algorithm that narrows hotspot positions by leveraging information across the genes that map to each hotspot (Supplementary Data 8 – 13). With this approach, we resolved the locations of 26 hotspots to regions containing three or fewer genes (a total of 58 genes; Figure 3A). Three hotspots contained exactly one gene (*GIS1*, *STB5*, and *MOT3*). We previously identified and experimentally confirmed the causal genes at several major hotspots (*MKT1*^27^, *HAP1*^14^, *IRA2*^16^, *GPA1*^29^, and the mating-type locus^14^), and these were all correctly localized by the algorithm, validating this fine-mapping strategy.

**Figure 3.**
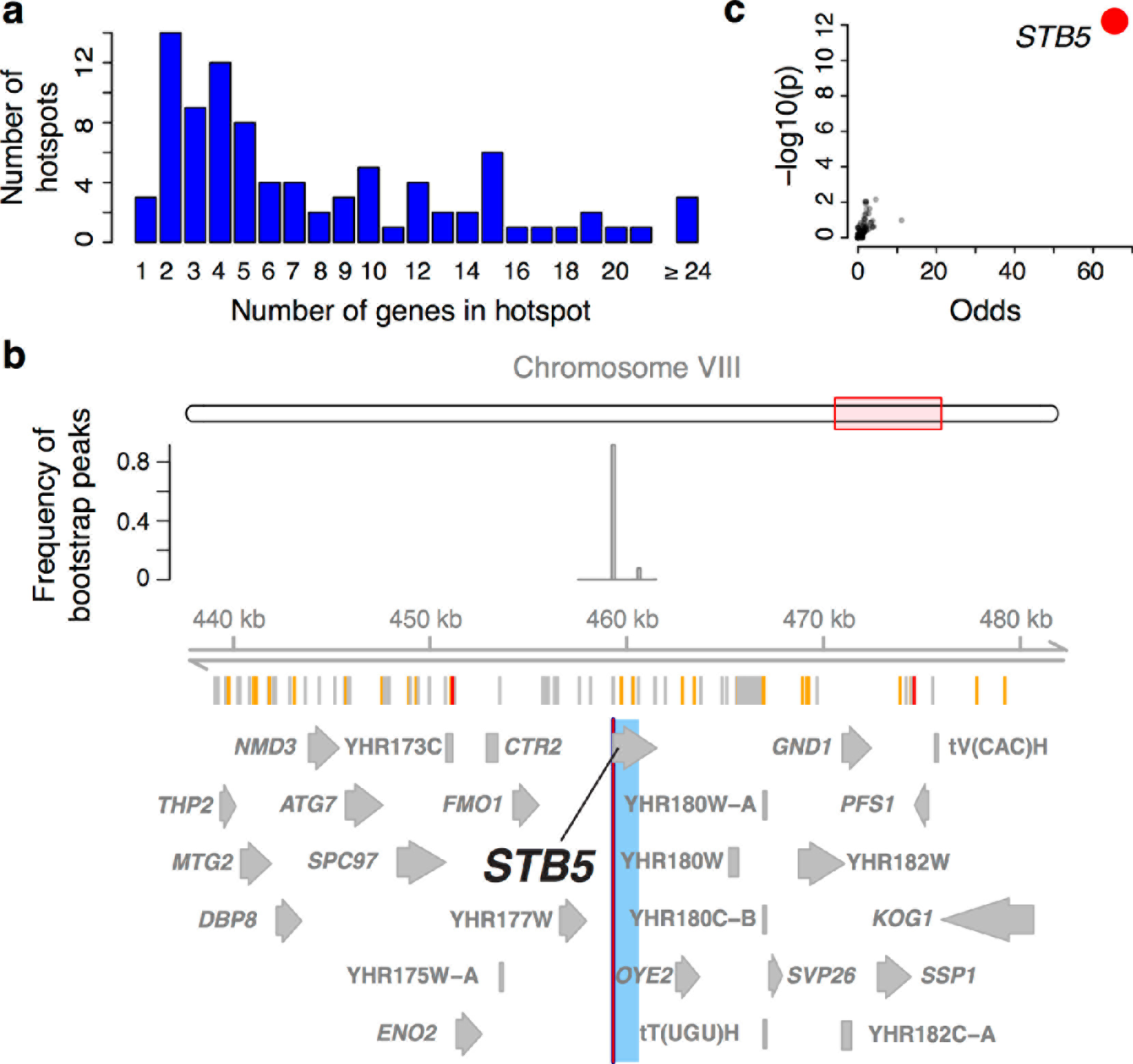
Genes located in hotspot regions. a) Histogram showing the number of genes located in the hotspot regions. b) A hotspot on chromosome VIII maps to the gene *STB5*. The figure shows, from top to bottom: a diagram showing the general region on the chromosome, the empirical frequency distribution of hotspot peak locations from 1,000 bootstrap samples (Methods), locations of BY/RM sequence variants (red: variants with “high” impact such as premature stop codons^67^; orange: “moderate” impact such as nonsynonymous variants; grey: “low” impact such as synonymous or intergenic variants), and gene locations. The light blue area shows the 95% confidence interval of the hotspot location as determined from the bootstraps. The red line shows the position of the most frequent bootstrap marker. c) Genes for which the BY allele at the *STB5* hotspot is linked to lower expression are enriched for *STB5* transcription factor (TF) binding sites in their promoter regions. The figure shows enrichment results for all annotated TFs (grey dots), with the strength of enrichment (odds ratio) on the x-axis vs. significance of the enrichment on the y-axis. The *STB5* result is highlighted in red.

The 58 genes at 26 high-resolution hotspots are highly enriched for the causal genes underlying the hotspots, making it possible to systematically study the functions of hotspot regulators. These genes were less likely to be essential or to have a human homolog than other yeast genes, but did not differ from other genes in their expression level or the number of physical or genetic interaction partners (Supplementary Table 5). The hotspot genes had highly significant enrichments for gene ontology (GO) terms related to transcriptional regulation (e.g. G0:0006357 “regulation of transcription from RNA polymerase II promoter”: 19 genes among the 58; 4 expected; p = 4e-9; Supplementary Figure 5; Supplementary Data 14), as well as weaker enrichments for terms related to response to nutrient levels (G0:0031669; “cellular response to nutrient levels”: 8 genes, 1 expected, p = 6e-6)

These analyses indicate that causal hotspot genes are disproportionately involved in transcriptional regulation, a signal that was not picked up in an earlier study with fewer, less-well-resolved hotspots^29^. For example, we fine-mapped a new hotspot that affected 382 genes to a single gene, the transcription factor *STB5* (Figure 3B). *STB5* is a transcriptional activator of multidrug resistance genes^30^. A previous analysis suggested reduced activity of the *STB5* BY allele compared to the RM allele^31^. Consistent with this observation, we found that the promoters of genes whose expression was lower in the presence of the *STB5* BY allele were strongly enriched for *STB5* binding sites (Figure 3C).

We fine-mapped a new hotspot on chromosome X to two genes (Supplementary Figure 6A). Of these, we deemed the transcription factor *CBF1* to be the more likely causal gene because the promoters of genes with higher expression in the presence of the BY allele at the hotspot locus were enriched for annotated^32^ *CBF1* binding sites (p = 2e-10; odds ratio = 5.4). *CBF1* regulates genes involved in sulfur metabolism^33,34^ and binds to centromeric DNA^33,35^ where it contributes to the production of centromeric transcripts^36^. However, the 307 genes that mapped to this locus were only weakly enriched for sulfur-related GO terms (e.g. GO:0072348 “sulfur compound transport”, p = 0.005) and showed no enrichment for centromere-related terms (Supplementary Data 12). Instead, the strongest enrichments were for terms related to the cell wall (e.g. GO:0071554 “cell wall organization or biogenesis”; p = 4e-10) and specifically the neck of budding daughter cells (GO:0005935 “cellular bud neck”; p = 0.0001). A potential role of *CBF1* in cell wall biology has been noted in the literature^34^ but has not been explored in detail. The strongest *trans* effect of this hotspot was on the gene *CAP1* (Supplementary Figure 6B), which is an annotated^32^ transcriptional target of *CBF1*. *CAP1* encodes a component of the capping protein heterodimer^37^, which localizes to the barbed ends of growing actin filaments where it prevents further actin filament growth^38^. In yeast, actin filaments play crucial roles in the transport of cell-wall material, vesicles, and organelles to the budding daughter cell^38^. Deletion of *CAP1* results in defective bud site selection in diploid yeast^39^. In our cross, higher expression of *CAP1* caused by the *CBF1* BY allele may increase actin filament blocking, altering the flow of cellular material during bud formation. In response, other genes involved in cell wall processes may adjust their expression, including strong *trans* targets of this hotspot such as *SLT2, CRH1*, and *DFG5* that are not annotated^40^ as direct targets of *CBF1*. This hotspot provides an example of an unexpected primary eQTL effect resulting in secondary effects, likely caused by changes in expression of the primary targets.

Hotspot genes can also influence mRNA levels more indirectly – for instance, by shaping the cellular response to external stimuli such as nutrient availability. For example, we fine-mapped a hotspot, which influenced 645 genes, to an interval on chromosome VIII containing six genes (Supplementary Figure 6C). One of these is *ERC1*, which encodes a transmembrane transporter. BY but not RM carries a frameshift in this gene, which removes the last two out of 12 predicted transmembrane helices of the protein^41^. This variant is known to reduce cell-to-cell variability (or “noise”) in the expression of a *MET17* gene tagged with green fluorescent protein^41^. We found that the BY allele at this hotspot reduced the expression of genes that are highly enriched for the GO category “methionine biosynthetic process” (GO:0009086, p = 2e-22; Supplementary Data 8 & 12). Thus, in addition to reducing *MET17* expression noise, the *ERC1* frameshift variant is linked to reduced mean expression levels of multiple genes in the methionine biosynthesis pathway (the *MET* regulon; Supplementary Figure 6D). While the precise compounds that are imported or exported by Erc1p are not known, the *ERC1* BY allele reduces cellular levels of S-Adenosylmethionine (SAM)^42^, a key component of methionine and cysteine amino acid metabolism^43^. The *ERC1* BY allele may down-regulate the *MET* regulon via its effects on SAM, triggering further transcriptional changes in hundreds of genes.

### Relationship of local eQTL and trans eQTL

Most known causal variants underlying yeast eQTL hotspots are coding; however, change in expression of a *trans*-acting factor by a local eQTL is another plausible causal mechanism^11^. We found that a higher proportion of hotspots contained genes with a local eQTL than expected by chance (p = 0.007; Figure 4A). The median effect size of the strongest local eQTL in these hotspots was larger than expected (p = 0.003). This enrichment is consistent with some hotspots being caused by local eQTL that alter the expression of a gene located at the hotspot position, which in turn leads to changes in the other transcript levels that map to the hotspot.

**Figure 4.**
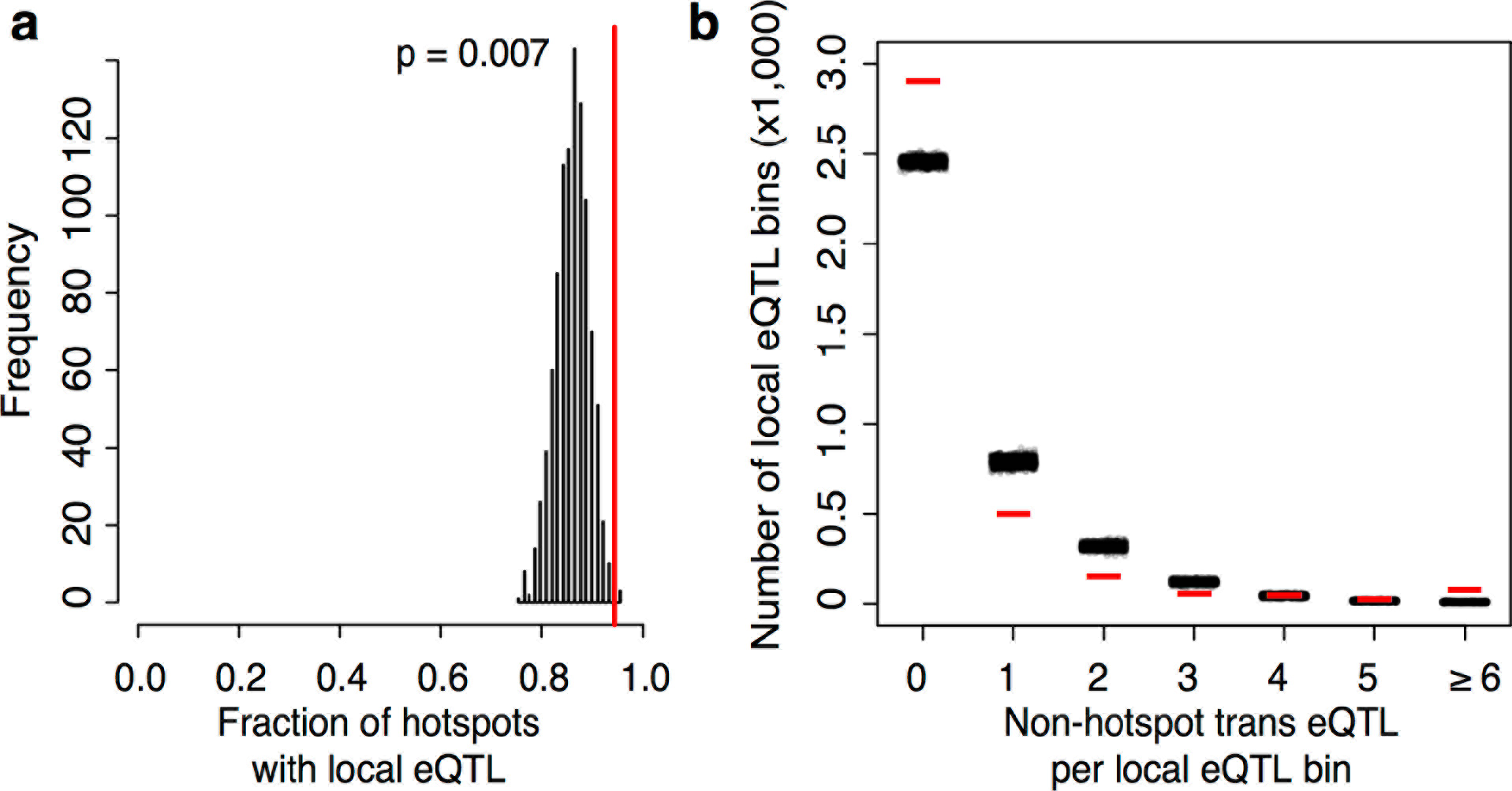
Relationship of local eQTL and distant eQTL hotspots. a) The fraction of hotspots that contain a genome-wide significant local eQTL. The black histogram shows the distribution observed in 1,000 random, size-matched regions of the genome. Because of the high number of local eQTL, most hotspots are expected to contain a local eQTL even by chance. The observed fraction (red line) still exceeds this random expectation. b) Distribution of *trans* eQTL at local eQTL in non-hotspot regions. The genome was divided into non-overlapping bins centered on local eQTL that did not overlap a hotspot. We counted the number of trans eQTL peaks in each bin. The figure shows the frequency (y-axis) of bins with a given number of trans eQTL (x-axis). The observed distribution is shown by red lines, and the distributions obtained in 1,000 randomizations of non-hotspot trans eQTL is show by clouds of black circles.

On the other hand, the majority of local eQTL (60%) did not overlap any of the hotspots. Evidently, the expression changes caused by these eQTL do not in turn lead to detectable *trans* effects on many unlinked genes, within the limits of our statistical power.

The density of *trans* eQTL in our data was sufficiently high, and their confidence intervals sufficiently wide, that any local eQTL overlapped at least two *trans* eQTL. However, this number is inflated due to the wide confidence intervals of the many weak *trans* eQTL. Using a more narrow definition of the *trans* signal, we focused on the peak markers of *trans* eQTL that did not overlap a hotspot. We divided the portion of the genome that did not overlap a hotspot into non-overlapping bins, each centered on a local eQTL. We then examined how many of the non-hotspot *trans* peaks fell into these bins (Figure 4B). The resulting distribution roughly matched the distribution expected if non-hotspot *trans* peaks were localized in the genome at random. To the extent that the distribution differed from random, we found an excess of bins with six or more *trans* eQTL (p < 0.001), likely reflecting undetected, weak hotspots. There was also an excess of bins with zero *trans* peaks (p < 0.001). This class comprised the great majority of the distribution (Figure 4B). The genetic architecture that is most consistent with these observations is one in which most local eQTL have no detectable *trans* consequences on the expression of other genes.

Even when the causal gene in a hotspot has a local eQTL, it does not automatically follow that this is the causal mechanism. For example, the *STB5*, *CBF1*, and *ERC1* hotspot genes each had a local eQTL. However, neither *STB5* nor *CBF1* showed allele-specific expression, and while there was weak allele-specific expression for *ERC1*, it was in the opposite direction of the local *ERC1* eQTL (Supplementary Data 7). Therefore, these local eQTL are unlikely to be caused by *cis*-acting variants.

*STB5*, *CBF1* and *ERC1* each carry protein-altering variants between BY and RM, including the known causal *ERC1* frameshift in BY. Altered protein activity due to these coding variants may be responsible for the many distant linkages to these three hotspots, and may also cause the observed local eQTL in *trans*, as previously shown for *AMN1*^23^. The Stb5p and Cbf1p transcription factors are both predicted to target their own respective promoters^32^, and their altered activity could influence their own expression. For the transmembrane transporter *ERC1*, the local eQTL may reflect *trans*-mediated feedback in which *ERC1* expression is increased in BY in an attempt to counter the reduced activity of the truncated *ERC1* BY allele. For each of these hotspots, it seems plausible that a change in protein function, rather than change in gene expression, underlies the hotspot.

### Genetics of mRNA vs. protein levels

The degree to which mRNA-based eQTL also affect the protein levels of their target genes is a fundamental open question^44–48^ that has been difficult to resolve as a consequence of low statistical power in eQTL and protein QTL (pQTL) studies. Low power is expected to lead to poor overlap between eQTL and pQTL solely as a result of high false-negative rates. We compared our eQTL to pQTL that we had identified earlier for 160 proteins using a powerful bulk segregant approach^18^ (Supplementary Data 15). These pQTL also clustered at hotspots, which broadly mirrored the mRNA hotspots identified here (Supplementary Figure 7). However, differences in hotspot architecture exist. For example, a hotspot on chromosome II shows strong pQTL effects^18^ but only weak effects on mRNA levels for the same genes, none of which rise to genome-wide significance.

In order to avoid downward bias in the overlap between eQTL and pQTL caused by false negatives (Supplementary Discussion 5), we focused on strong QTL in each dataset and asked if they overlapped a significant QTL in the other dataset. Of the 236 strongest eQTL (variance explained ≥ 3.5%; ≥ 99% power to detect), only 47% (111) overlapped a pQTL for the same gene. Of the 218 strongest pQTL (LOD ≥ 15), 50% (108) overlapped an eQTL for the same gene. Thus, even with high power and strong QTL effects, agreement between eQTL and pQTL was imperfect. Strong eQTL without a pQTL clustered primarily at the *HAP1* and *MKT1* hotspots (Supplementary Table 6; Supplementary Figure 8B). These two hotspots also showed the clearest examples of overlapping eQTL and pQTL with opposite direction of effect on the same genes (Supplementary Table 7; Supplementary Figure 8A). Thus, while these hotspots influence both mRNA and protein levels of many genes, their effects on mRNA vs. protein levels of a given gene can be quite different. Strong pQTL without an eQTL were more widely distributed across the genome (Supplementary Table 8; Supplementary Figure 8C).

### Detection of non-additive eQTL interactions from a genome-wide search

The contribution of non-additive or “epistatic” genetic interactions to trait variation is a topic of ongoing debate^49–51^. In particular, demonstration of non-additive effects on human gene expression has been challenging^52–57^. Although clear examples of epistasis have been revealed for yeast gene expression^58,59^, the limited power of earlier studies had necessitated targeted search strategies rather than a full genome-by-genome scan.

We reasoned that the high power of our current dataset should permit a more unbiased view of the contribution of epistasis to mRNA expression variation, and carried out a genome-by-genome scan for non-additive interaction effects on the expression levels of all genes. We detected 387 eQTL-eQTL interactions (FDR = 10%; Figure 5A; Supplementary Data 16). To our knowledge, this is the first unequivocal identification of eQTL interactions from an unbiased genome-by-genome scan. Targeted scans with a reduced multiple testing burden identified larger numbers of interacting pairs of loci: a total of 784 from a scan for interactions between genome-wide significant additive eQTL and the genome, and a total of 1,464 for interactions between significant additive eQTL. As previously reported, the interactions occurred primarily between pairs of loci in which one or both loci had a genome-wide significant additive effect. In particular, many pairs involved interactions between strong eQTL at major *trans* hotspots^60^, or interactions between strong local eQTL and distant eQTL at hotspots (Figure 5A).

**Figure 5.**
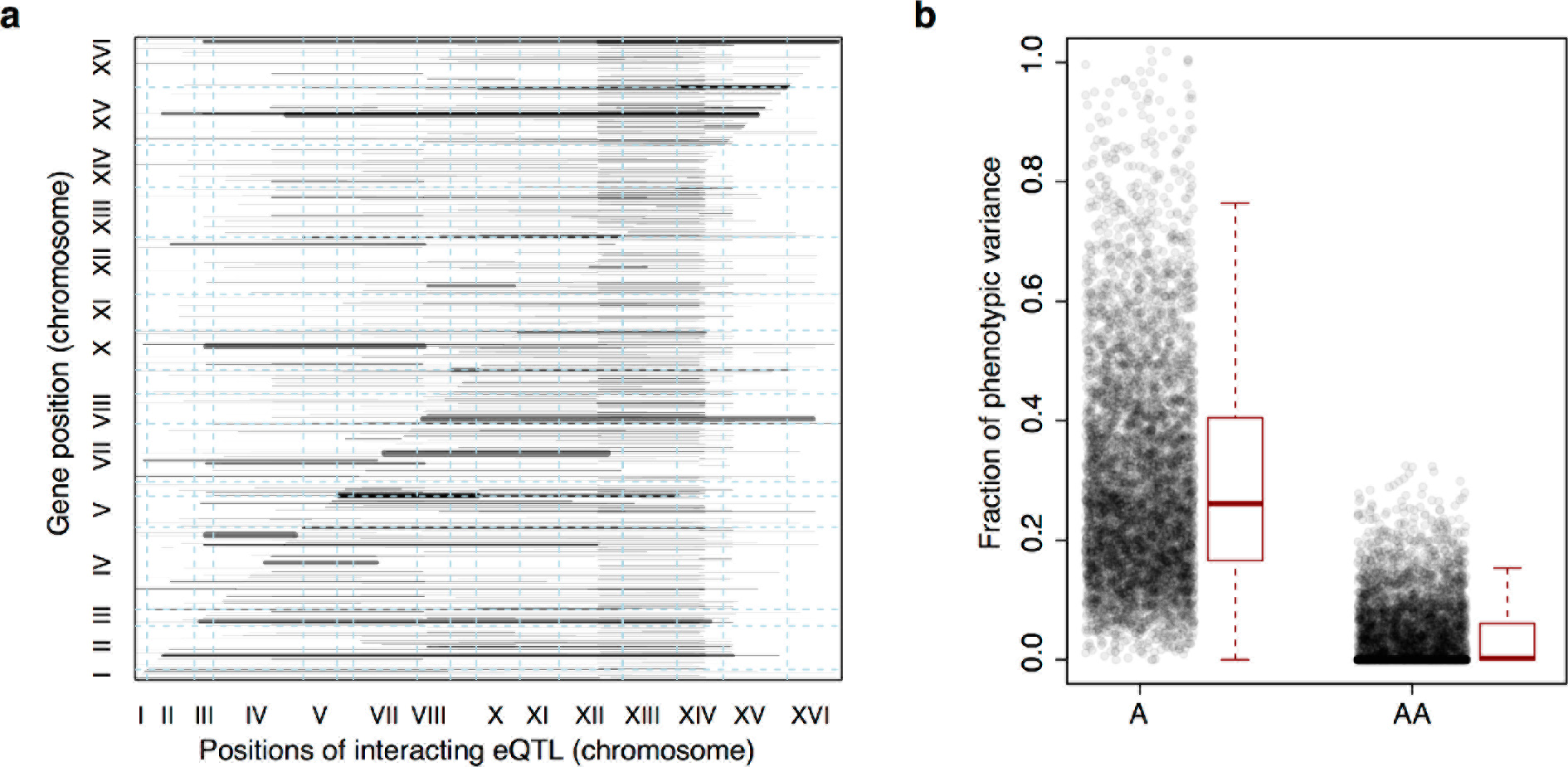
Non-additive interactions between eQTL. a) Map of detected interactions between pairs of eQTL. The position of the affected gene is shown on the y-axis. The x-axis shows the locations of the two interacting eQTL in each pair, which are connected by horizontal lines. Line width scales as a function of the strength of the interaction term. Only pairs with a significant interaction term detected in a genome wide search (all genome positions by all genome positions for all expressed genes) are shown. eQTL pairs often involve a local eQTL (visible as line endpoints falling along the diagonal) and distant eQTL hotspots (visible as line endpoints forming vertical bands). b) Boxplots of the fraction of phenotypic variance (y-axis) explained by genetic variation as captured by genome-wide relatedness in an additive (“A”, left), and interactive (“AA”, right) model.

We quantified the fraction of variation in gene expression that is contributed by epistatic interactions. Pairwise interactions typically explained about 1/10th as much expression variance as did additive loci (Figure 5B). Thus, genetic interactions contributed only a small minority of trait variance for gene expression levels, which is consistent with what we previously reported for organism-level traits^21^.

### Conclusion

The high power of our study allowed us to identify genome-wide significant eQTL that jointly explain over 70% of gene expression heritability. The completeness of our eQTL catalog allowed us to examine the contribution of *trans*-acting regulatory variation in much greater detail than previously possible. We showed directly that *trans*-acting eQTL form the predominant source of expression variation in a yeast cross, in agreement with indirect genome-wide estimates in humans. We also showed that the vast majority of *trans*-eQTL were concentrated at a limited number of hotspot regions that are inferred to harbor variants with widespread effects on the expression of other genes. Indeed, the strongest hotspots affected the expression of most genes in the genome. We showed that hotspots were enriched for genes involved in transcriptional regulation. The minority of eQTL that fell outside the statistically defined hotspots also clustered more than expected by chance, and we saw little evidence of isolated *trans*-acting loci that affect the expression of one or a few genes. The limited number of human *trans*-eQTL discovered to date also tended to influence the expression of multiple genes^3,6–13^, suggesting that a similar hotspot-dominated architecture underlies human expression variation and will be uncovered in better powered studies.

The recently proposed omnigenic model^61^ for the genetic basis of complex trait variation posits that gene regulatory networks are sufficiently densely connected that the change in expression of any one gene, caused by a local eQTL, will “percolate” through the network and alter the expression of all other genes. The hotspot loci we described here offer evidence that some regulatory variants can indeed have widespread effects on the transcriptome, in some cases altering the expression of the majority of genes in the genome through precisely the combination of strong direct effects on “core” genes in specific pathways and weak indirect effects on other “peripheral” genes envisioned in the omnigenic model.

On the other hand, although we detected local eQTL for most genes in our cross, the majority of these had no detectable *trans* effects on the expression of other genes, within the limits of our statistical power. Given that our study had sufficient power to detect weak indirect effects of *trans*-eQTL hotspots, we believe that most local eQTL indeed have no meaningful downstream consequences for gene expression, and, by extension are unlikely to contribute to variation in complex traits. Consistent with this conclusion, modest expression changes for dozens of yeast genes have been found to result in minimal fitness effects^62^. These results argue against the simplest form of the omnigenic model, in which a variant that changes the expression of any one gene has meaningful effects on every other gene. Instead, we observed that *trans*-eQTL effects preferentially arise from variation in certain classes of genes, and may be caused by coding as well as regulatory variants. Given the crucial importance of regulatory variation for many complex traits, the organismal consequences of expression changes caused by different types of eQTL remain a key area for further research.

## Methods

Unless otherwise specified, all computational analyses were performed in R.

### Yeast growth

We used 1,012 meiotic segregants previously generated^15^ from a cross between the prototrophic yeast laboratory strain BY (*MATa;* derived from a cross between BY4716 and BY4700) and the prototrophic vineyard strain RM (*MATα hoΔ::*hphMX4 *flo8*Δ::natMX4 *AMN1-BY*; derived from RM11-1a). The segregants were grouped according to their previously measured^15^ endpoint colony radius on YNB agar plates into groups of 96. The strains in each group were rearranged from existing stock plates into a total of 13 96-well plates in YNB medium, grown to saturation, and frozen as glycerol stocks for later growth. Within each group of 96, strain locations in the 96-well plate were selected at random. Culture and liquid handling was performed on a BioMek FXP instrument or with multichannel pipettes in 96-well format.

Our strategy of batching segregants according to their growth on YNB ensures that each 96-well plate contains segregants that grow at comparable rates. This facilitates growing all segregants on a plate such that they reach a similar optical density at 600 nm (OD) at the same time. Our batching strategy produces experimental batches that are correlated with growth rates. Because we statistically removed variation among experimental batches prior to eQTL mapping (see below), this design reduces our ability to compare variation in growth rates with variation in gene expression. We deemed this an acceptable trade-off because it considerably simplified handling >1,000 samples in a systematic fashion. We processed the batches in a randomized order with respect to their growth rate to avoid confounding processing date with faster or slower growth.

We used the rearranged stock plates to inoculate growth cultures in 1 ml YNB medium (recipe for 1 L: 6.7g yeast nitrogen base with ammonium sulfate and without amino acids; 900 ml H_2_O; autoclave; add 100 mL of separately autoclaved 20% glucose solution) in 2 mL deep well plates sealed with Breathe-Easy membranes (Sigma Aldrich), and grew the cultures to saturation on Eppendorf MixMate instruments situated in a 30°C incubator and set to 1100 rounds per minute (rpm). We set the saturated cultures back to OD = 0.05 in 1 mL YNB in a fresh deep well plate and continued growth at 30°C. We monitored OD during growth by splitting out 100 ul of culture every other hour, measuring OD on a Synergy 2 plate reader (BioTek) and returning the 100 ul used for measuring OD to the deep-well culture plate. We increased the frequency of measurements as cultures approached OD = 0.4.

Once average OD in the plate reached 0.4, we transferred the cultures to sterile Norgen nylon filter plates (#40008) situated on a vacuum manifold. We applied vacuum to remove all growth medium, sealed with aluminum foil seals, and flash froze the entire plate in liquid N_2_. The frozen plates were placed on a standard 96-well plate to protect their bottom, wrapped with parafilm, and stored at −80°C until RNA extraction. Note that this procedure provided us with OD measurements up the exact time point at which cells were harvested.

### RNA extraction

We used Dynabeads mRNA purification kits (Ambion / Thermo Fisher) to directly isolate mRNA from cell lysates. To perform the RNA extractions on the BioMek robot, we prepared excess lysis/binding and Wash buffers that permitted the use liquid reservoirs with volumes that exceed that provided in the kits. These buffers were prepared as specified in the Dynabeads kit protocol:

*Lysis*/*Binding Buffer:*

100 mM Tris-HCl pH 7.5
500 mM LiCl
10 mM EDTA, pH 8
1% LiDS
5 mM dithiothreitol (DTT)

*Washing Buffer A:*

10 mM Tris-HCl, pH 7.5
0.15 M LiCl
1 mM EDTA
0.1% LiDS

*Washing Buffer B:*

10 mM Tris-HCl, pH 7.5
0. 15 M LiCl
1 mM EDTA

We filled the wells of an Axygen 1.1 mL plate (P-DW-11-C-S) with about 250 *μ*l acid washed 425-600 *μ*m beads (Sigma G8722). We added 700 *μ*L lysis buffer to our frozen cell plates, pipetted up and down to resuspend the cells, and applied them to the glass beads in the Axygen plate. The Axygen plate was tightly sealed with an Axymat rubber plate seal (AM-2ML-RD-S), and ground for 10 cycles on a plate-based mini bead beater (Biospec). Each cycle consisted of 1 minute beating followed by 1 minute on ice.

We centrifuged the plate for 4 minutes at 3,000 rpm to separate glass beads and cell debris from the lysate. We pipetted two aliquots of 200 *μ*L of lysate supernatant into two 96-well PCR plates for a total of 400 *μ*L lysate. These plates were sealed, and the RNA melted for 2 minutes at 65°C in a thermocycler. We implemented a BioMek-assisted procedure to perform the Dynabead protocol with two mRNA enrichment steps. We did not quantify the resulting 11 *μ*L of mRNA and simply used the entire mRNA for reverse transcription and sequencing library preparation. While piloting this procedure, we obtained typical yields of ~30 ng / *μ*L and excellent RNA quality as judged by visualization on 1.1% agarose gels stained with ethidium bromide. Ribosomal RNA bands were clearly visible in crude lysate, less visible after the first mRNA enrichment, and absent after the 2^nd^ mRNA enrichment step. After the 2^nd^ mRNA enrichment, mRNA was clearly visible on the gel, with no visible RNA degradation.

### RNA sequencing library construction & sequencing

We performed reverse transcription and sequencing library preparation using the Kapa Stranded mRNA-Seq Kit (KK8420/21). This kit usually begins by enriching mRNA from total RNA. Because we had already performed mRNA enrichment, we used our entire mRNA as input and began at the RNA fragmentation step by adding 11 *μ*L of “KAPA fragment, prime and elute buffer” to our 11 *μ*L of mRNA. RNA fragmentation was performed on a thermocycler for 6 minutes at 94°C.

The remaining procedure was performed as specified in the Kapa kit manual. Briefly, the fragmented RNA is randomly primed and used for 1^st^ strand cDNA synthesis, 2^nd^ strand synthesis and marking with dUTP, A-tailing of the double-stranded cDNA, adapter ligation, and PCR for 12 cycles. The dUTP marked 2^nd^ strand is not amplified in PCR, resulting in strand-specific libraries. We used custom designed Truseq-compatible indexing adapters (IDT) to allow multiplexing all 96 samples per batch. Prior to use, the two types of Truseq adapters were annealed (2 minutes at 97°C; 72 steps of 1 minute at 1°C decreasing temperature; 5 minutes at 25°C) to generate forked adapters that can be ligated to the A-tailed cDNA. We did not pool samples between batches.

Sequencing libraries were quantified by combining 1 *μ*L of library with 100 *μ*L of Qubit High Sensitivity dsDNA reagent in 96-well plates with black bottom and wells, and reading fluorescence (excitation 485nm, emission 528 nm) on the Synergy 2 plate reader. We calculated library concentrations by comparing to a standard series obtained by diluting the standard solutions included in the Qubit quantification kit. Standards were measured in triplicate on each library plate. We pooled the libraries in each group to equal molarity and used qPCR (KAPA Biosystems #KK4854) on the pool to obtain the molarity for loading on the sequencer. Gel extraction was not necessary because the RNA fragmentation and bead clean-up that are part of the Kapa protocol resulted in library fragments of the desired size of 200 - 400 bp.

Sequencing was performed for 100 bp single end on Illumina HiSeq 2500 instruments at the UCLA BSRC sequencing core for two lanes per batch, for 26 total lanes. On average, we obtained approximately 3 million reads per sample.

### Sequence processing & gene expression quantitation

Adapter sequences were trimmed using trimmomatic^63^. Reads were pseudoaligned to the 6,713 annotated yeast ORF coding sequences from Ensembl build R64-1-1 using kallisto v.43.0^64^. Kallisto was run in strand-specific mode with parameters −l 150 and –s 8. For each transcript, we computed transcripts per million reads (TPM) as a measure of expression and used log2(TPM + 0.5) for downstream analysis. Segregants with fewer than one million reads were removed from downstream analysis, and 1,012 segregants passed this filter. We removed 993 invariant transcripts with identical expression across all segregants or with log2(TPM + 0.5) less than 1 in 50% or more of the segregants. Our final dataset included 5,720 transcripts, which were used for downstream analyses. These transcripts cover 5,506 of 5,971 open reading frames annotated as ‘verified’ or ‘uncharacterized’ in the yeast genome^40^.

### Growth Rate Covariate

Unless otherwise specified, all remaining analyses were conducted in R (www.r-project.org). Based on the OD measurements collected during growth prior to harvesting, growth rates were calculated for each segregant using the R package grofit and the function gcFitSpline^65^. The difference between the maximum and minimum OD was recorded for each culture and used as a covariate for downstream analysis.

### Sequence variants

Our BY and RM parent strains had earlier been sequenced to very high depth (>200-fold coverage of the genome), and GATK^66^ used to identify 48,254 sequence variants between them. These variants (irrespective of whether or not they are part of our marker map) were screened for potential functional impact using the Ensembl Variant Effect Predictor^67^.

The segregant genotyping is described in Bloom 2013^15^ and Bloom 2015^21^. The 1,012 segregants used for this study were genotyped at 42,052 highly reliable markers, which are a subset of the total 48,254 sequence differences between BY and RM. Sets of markers that were in perfect linkage disequilibrium (i.e. markers never separated by recombination) among the 1,012 segregants were collapsed to one marker. Our final linkage map comprised 11,530 unique markers.

### Heritability

A variance component model was used to estimate additive heritability. First, gene expression measurements were corrected for batch covariates and the growth measurement covariate described above using a linear model for each gene

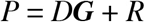

where *P* is a vector of log_2_(TPM + 0.5) measurements for *n* segregants for that gene. *D* is a vector of estimated fixed effect coefficients for technical covariates. ***G*** is a matrix of *n* total segregants by *m* technical covariates. Technical covariates included experimental batch and the growth rate covariate described above. The vector of residuals is denoted as *R*. *R* contains expression phenotypes corrected for batch effect and growth covariate.

We fit the variance component model

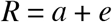

where *a* is the vector of additive genetic effects, and the residual error is denoted by *e*. The distributions of these effects are assumed to be normal with mean zero and variance-covariance as follows:

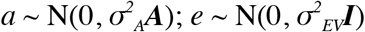

Here, ***A*** is the additive relatedness matrix - the fraction of the genome shared between pairs of segregants. ***A*** was calculated using the ‘A.mat’ function in the rrBLUP R package^68^. 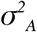 is the additive genetic variance captured by markers. 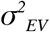 is the error variance and ***I*** is the identity matrix. Additive heritability was estimated using custom code adapted from Kang et al^69^.

Although our heritability estimates are lower bounds due to counting noise in the number of sequencing reads per gene, downsampling of reads suggested that additional sequencing would increase heritability for most genes by at most a few percent (Supplementary Figure 1).

Additionally, we fit a model to estimate the relative contribution of pairwise interactions with

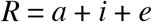

where 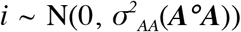 and ***A**°**A*** is the Hadamard (entry-wise) product of **A**, which can be interpreted as the fraction of pairs of markers shared between pairs of segregants. 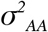 is the interaction genetic variance captured by all pairwise combinations of markers. The other terms are the same as in the additive-only model. The result of ~1/10 as much variance arising from interactions relative to additive loci is based on the ratio of the average of the ***A**°**A*** term across genes to the average of the ***A*** term across genes. When we instead calculated this variance ratio for each gene, we found that the mean across genes was greatly inflated by a few extreme outliers, while the median was very low (less than 1/100) because almost half of the genes had an estimate of zero for the ***A**°**A*** term.

### Gene annotations and features

Gene positions were extracted from Ensembl^70^ (www.ensembl.org) build 83. Various analyses throughout the paper made use of a range of gene-specific features, factors and covariates: 1) Total variance in expression was calculated as the sum of the additive and residual variance components obtained in our heritability estimates. 2) Expression level was calculated as the mean log2(TPM) across segregants, 3) Gene essentiality was coded as a binary factor and obtained from SGD^40^ (www.yeastgenome.org) by searching for genes whose SGD deletion phenotype contained the term “inviable”. 4) dNdS values were obtained from Supplementary Table S4 in Wall et al., 2005^71^. 5) The number of protein-protein interactions was obtained from SGD by downloading all “physical” interactions between genes and counting their number per gene. 6) Synthetic genetic interactions were extracted from data from Costanzo et al., 2016 which provides genetic interaction data for pairwise gene deletions or disruptions between nearly all essential (E) and nonessential (N) genes^72^. Specifically, we downloaded the “NxN”, “NxE”, and “ExE” raw genetic interaction datasets from http://thecellmap.org/costanzo2016/, combined them into one table, and extracted the lowest interaction p-value for each gene pair. We restricted this set using the “strict” definition from Costanzo et al.^72^ and kept only pairs with interaction p-value < 0.05 and interaction strength (epsilon) > 0.16 or < −0.12. For each gene, we counted how many genes showed a genetic interaction at these thresholds and used this as our measure of synthetic genetic interactions. Using the “lenient” or “intermediate” definitions did not alter our conclusions. 7) We defined whether or not a gene is a transcription factor by downloading from SGD all genes annotated to the GO term GO: 0003700 “transcription_factor_activity_sequencespecific_DNA_binding” and its child GO terms. 8) As a proxy for deep evolutionary conservation, we extracted from Ensembl biomart whether or not a gene has a human homolog.

Gene ontology (GO) associations for each gene were downloaded from the Gene Ontology Consortium (geneontology.org) on February 16, 2016. We used paralogy information downloaded from the yeast gene order browser^73^ (http://ygob.ucd.ie/).

### Characteristics of genes with high or low heritability

We tested for gene features associated with the degree of heritability by multiple linear regression. This regression modeled heritability as the dependent variable and the various gene features as predictor variables. We used the ‘summary’ and ‘lm’ functions, and the ‘car’ package in R to perform Type III sum-of-squares ANOVA. This analysis tests for the influence of each feature by dropping it from a full model that includes all other terms, and asking whether this results in a significantly worse fit as judged by F-statistics. The analysis controls for correlations among predictor variables and reports marginal associations only if they are significant over all other terms. We did not include interaction terms among predictor variables.

### Gene ontology enrichment analyses

We tested for GO enrichments using the R package topGO^74^. For analyses in which genes were classified as “interesting” or not (e.g. whether a gene has heritability ≥ 90%, or whether it is located in a hotspot), we used the Fisher test for enrichment. When using a quantitative gene score as the measure of interest (e.g. the heritability), we used the one-sided t.test implemented in topGO. We used the ‘classic’ scoring method^75^, i.e. we did not adjust the enrichments for significance of child GO terms.

### eQTL mapping

We^21^ and others^76,77^ have previously noticed that power and precision of QTL mapping on a given chromosome can be increased by controlling for genetic contributions that arise from the other chromosomes in the genome. Our eQTL mapping strategy controls for genomic background in two ways. For each gene, we identified large genetic effects segregating on other chromosomes and included them as covariates while mapping on a given chromosome. We also corrected for any additional polygenic additive background signal on other chromosomes. Then, for each gene we used a forward stepwise procedure to map eQTL with a false discovery rate procedure. Below we describe our algorithm in greater detail. Throughout, we use the terms “eQTL” and “linkage” interchangeably.

#### Identification of large background genetic effects

In the process of eQTL mapping on a given chromosome, we wanted to control for the genetic contributions from the remainder of the genome. Although this background control is sometimes done using a polygenic model with a random effect that captures overall relatedness across the genome^76^ (see below), large individual eQTL effects may not be adequately accounted for by one genome-wide relatedness matrix. Therefore, we performed the following procedure to identify large genetic effects. Our goal at this stage was not to formally identify these large effects as eQTL, but to perform a simple scan for large effects that can be included as covariates to control for their effects while mapping on a given chromosome. As our algorithm progresses along each chromosome, these large background effects will eventually be detected as formal eQTL.

1. Gene expression measurements were corrected for batch covariates and the growth covariate using a linear model

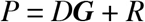

where *P* is log2(TPM + 0.5), *D* is the vector of estimated fixed effect coefficients for technical covariates (batch and growth), and ***G*** is a matrix of *n* total segregants by *m* technical covariates as described in the section on Heritability above. The residual vector is denoted as *R* and contains the corrected expression measurements.
2. We sought to identify a set of markers linked to large-to-moderate effect QTL. For each gene and for each chromosome, we calculated logarithm of the odds (LOD) scores between corrected expression levels *R* and the marker genotypes on that chromosome as

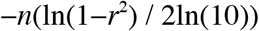

where *n* is the number of segregants with genotypes and phenotypes, and *r* is the Pearson correlation coefficient between segregant genotypes and *R*. The marker with the largest LOD per chromosome was added to a matrix, ***Z***, if it had LOD > 3.5.
3. For each gene we calculated

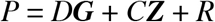

where *C* is a vector of genotype effects. This step controls for technical covariates as above, but additionally controls for large effects included in ***Z***. It results in a new vector of residual expression levels *R*. We repeated steps 2 and 3 twice with this new *R*, appending additional markers to ***Z*** for each gene and each chromosome if they passed the threshold of LOD > 3.5. The goal of this repeated search for large effect eQTL was to control for large effect loci that were detected only after expression values were corrected for the effects of previously identified large effect loci. For example, a given chromosome may harbor several large effect QTL, and repeated runs of steps 2 and 3 ensure that such loci are captured. At the end of this procedure, we had a matrix ***Z*** of up to three markers per chromosome per gene that were linked to large-to-moderate effect loci.

#### Correcting gene expression measurements for large background effects and additive polygenic background for all chromosomes except the chromosome of interest

For each chromosome of interest and for each gene expression trait we calculated

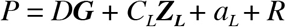

*C_L_* and ***Z***_*L*_ are the background eQTL effects identified from the procedure above that are not located on the chromosome of interest. 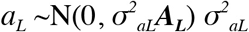 is the additive genetic variance from all chromosomes excluding the chromosome of interest. ***A_L_*** was calculated using the ‘A.mat’ function in the rrBLUP package using a genetic relatedness matrix that excludes markers from the chromosome of interest. The goal of this step was to obtain expression phenotypes *R* that can be used to scan for eQTL on a given chromosome by correcting for sources of variation that do not arise from that chromosome: batch and growth effects, large effects on other chromosomes, and a polygenic term accounting for any additional genetic contributions arising from other chromosomes.

#### Mapping additive eQTL

We mapped additive eQTL using a forward stepwise procedure. For each chromosome and for each gene we tested for linkage at each maker on the given chromosome with residual expression values *R* (calculated above) using the formula in Step 2 of “Identification of large background genetic effects”. We recorded the location and LOD score of the marker with the highest LOD score. To decide if this marker should be included as a QTL in the model, we used a permutation-based FDR criterion of 5%.

FDR was calculated as the ratio of the number of genes expected by chance to show a maximum LOD score greater than a particular LOD threshold vs. the number of genes observed in the real data with a maximum LOD score greater than that threshold, for a series of LOD thresholds ranging from 1.5 to 9 with 151 equal-sized steps of 0.05. The number of genes expected by chance was calculated by permuting *R* relative to segregant genotypes, calculating LOD scores for all genes across the chromosome and recording the maximum LOD score for each gene. In each run of the permutations, the permutation ordering was the same across all genes. We repeated this permutation procedure 1,000 times. Then, for each of the 151 LOD thresholds, we calculated the average number of genes with maximum LOD greater than the given threshold across the 1,000 permutations. We found the lowest of the 151 LOD thresholds at which the ratio of the number of expected genes with an eQTL of at least this threshold to the number of observed genes of at least this threshold was < 5%. This LOD threshold was used as a criterion for declaring a given QTL in the real data as significant.

For all genes with a significant linkage (FDR <5%), the peak marker was added to the linear model for that gene as *X*:

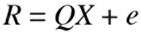

Genes without a significant linkage were excluded from additional testing on that chromosome. For the set of genes with a significant linkage, we repeated the procedure above by replacing *R* with *e*. The procedure was repeated for each chromosome until no genes had additional significant linkages.

### Cross-validation

The amount of additive variance explained by detected QTL was estimated using crossvalidation. Segregants were grouped based on the batches used for RNA and library preparation. Each batch of segregants was left out of the procedure one at a time. The eQTL mapping procedure was performed for all the other batches. For the QTL markers detected in this training set, the amount of variance explained by the joint model of all significant QTL markers was estimated in the held out batch.

### eQTL confidence intervals

eQTL confidence intervals were calculated as 1.5 LOD drops. We extended the eQTL location confidence intervals to include all markers in perfect LD with the markers used in eQTL detection (marker correlation = 1).

### Hotspot identification

We devised an algorithm with the goal of identifying a set of eQTL hotspots by combining information across genetically correlated transcripts and, most importantly, using co-localizing trans-eQTL to better narrow hotspot confidence intervals. The algorithm has three major steps. First, we control for unmodeled factors affecting gene expression that may obscure hotspot detection and localization. Second, we use a multivariate statistic to identify eQTL hotspots. Finally, we use a bootstrap procedure to delineate confidence intervals for hotspot location. We describe the steps in greater detail below.

#### Identification of unmodeled factors affecting gene expression

1. For each gene with at least one statistically significant eQTL we fit a linear model

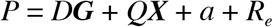

where *P* is log2(TPM + 0.5), *D* is the vector of estimated fixed effect coefficients for technical covariates, and ***G*** is a matrix of *n* total segregants by *m* technical covariates as described in the section on Heritability. ***X*** is a matrix of the statistically significant additive QTL peak markers for the gene, *Q* is a column vector of QTL effects, and *a* is random effect for the contribution of polygenic background to additive variance, where 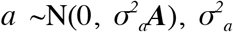 is the additive genetic variance from all markers and ***A*** is constructed as described above in the section on calculating heritability. *R_e_* is the residual expression level after the contributions of all additive influences on gene expression variation in the model have been removed.
2. For each gene, *R_e_* was scaled to have mean 0 and variance 1. *R_e_* for each gene was concatenated to form the columns of the matrix ***R***.
3. We calculated the singular value decomposition (SVD) of ***R***. We also calculated the SVD after each column of ***R*** was individually permuted. We visually inspected a Scree plot for both decompositions and observed that the top 20 eigenvectors explained more variance than expected by chance.
4. The top 20 eigenvectors were appended to the matrix ***G*** of covariates. These eigenvectors capture systematic expression variation that is shared across genes but that does not arise from any known technical covariate, identifiable eQTL, or additive genomic background as captured by our markers. These components presumably reflect undetected sources of variation from unmodeled experimental factors or non-additive genetic factors shared across genes. Their inclusion as covariates in ***G*** below follows from the same logic that motivates SVA^78^; we want to account for unmodeled factors that are contributing to gene expression variation to increase power to map and resolve eQTL hotspots.

#### Correcting gene expression for technical covariates, detected additive QTL on other chromosomes, and unmodeled factors

For each chromosome, we extracted all genes that have a significant linkage to that chromosome but do not physically reside on that chromosome (i.e., all genes that had a statistically significant trans acting eQTL on the given chromosome) and fit the linear model

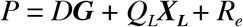

This model is similar to that in Step 1, “identification of unmodeled factors affecting gene expression”, except that the matrix ***G*** is extended to include the 20 eigenvectors corresponding to unmodeled factors, and ***X_L_*** only contains statistically significant additive eQTL that do not reside on the chromosome of interest. Thus, *R_e_* is corrected for all influences on expression that are not relevant to mapping hotspots on the given chromosome, but retains all signal that does arise from the chromosome of interest.

For each gene, *R_e_* was scaled to have mean 0 and variance 1. *R*_*e*_ for each gene was concatenated to form the columns of the matrix ***R***.

#### Dimensionality reduction of R

Our algorithm for fine-mapping eQTL hotspots makes use of a multivariate statistic (see section “multitrait mapping to localize eQTL hotspots”). We observed that on many chromosomes, the number of genes with *trans* linkages exceeded our total sample size. Because the multivariate statistic is not defined in this case, we reduced the dimensionality of ***R*** using SVD. The top *m* eigenvectors with corresponding eigenvalues greater than those observed from an SVD on permuted data were retained as matrix ***L***. The entries of ***L*** can be interpreted as weighted linear combinations of the expression levels from individual genes, from which all sources of variation that do not arise from the chromosomes of interest have been eliminated. These combinations capture the majority of additive genetic influences on the given chromosome that are shared across genes (i.e. the effects of hotspots on multiple genes), and serve as the input to our hotspot detection algorithm.

#### Multitrait mapping to localize eQTL hotspots

We computed

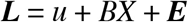

where *u* is a vector of means for each of the columns of ***L***, *B* is a vector of coefficients for the effects of *X* on each of the columns of ***L***, *X* is the genotype vector for each marker on the target chromosome (the model is fit one marker at a time), and ***E*** is a matrix of residuals. Next, we calculated the residual sum of squares matrix ***RSS*** as ***E`E*** and LOD scores as (n/2)log_10_(*|**RSS_0_**|/|**RSS**|*), where |**RSS**| denotes the determinant of the ***RSS*** matrix and ***RSS_0_*** is the residual sum of squares matrix for the null model with no QTL effect^79,80^. This LOD score reflects linkage of genetic markers with the joint expression of the eigenvectors in ***L***. To distinguish significant linkages affecting multiple genes from eQTL for individual genes, we refer to them as jointQTL (jQTL) in the remainder of this method section.

We fit the equation above 100 times with permutations of phenotype to genotype for matrix ***L***. The 99% quantile of the maximum observed LOD score per permutation was used to decide whether the maximum multivariate LOD was significant. If it was significant, then the effect of that marker was subtracted from ***L*** and the procedure repeated until no more jQTL were detected.

#### Refining jQTL locations and test for hotspots that are better modeled as two neighboring hotspots

We sought to better refine the location of the jQTL given all other jQTL on that chromosome^81^. We collected the statistically significant peak markers for the jQTL identified above. Each peak marker was dropped from a joint model one at a time and the peak position was recomputed. Peaks that moved by more than 75 kb at this step were removed from the model. For all peaks with LOD > 200 we tested the possibility that the multivariate peak was a “ghost” jQTL, i.e. a jQTL that appears to be localized between two or more true jQTL that are very closely linked and that influence a similar set of genes. We fit a model where the one significant jQTL was replaced by a two-jQTL model, with the added criterion that the two jQTL could not be within 10 markers on either side of the original peak. Permutations for this two-locus model were performed as in “multitrait mapping to localize eQTL hotspots”, with the relevant test and null statistics being the difference between the fit of the best two-locus model and the best one-locus model. Single jQTL peaks were replaced with the best two-locus model if the observed LOD was more than 10 greater than the 99% quantile LOD for the permuted data. Through the combination of the steps above, this procedure identified significant jQTL that correspond to hotspots that influence the expression of multiple genes.

#### Bootstrap resampling to identify confidence intervals for jQTL location

For each detected jQTL, segregants were sampled with replacement 1,000 times. For each sample, we refit

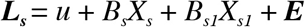

where *s* indicates a bootstrap resample of the rows of a matrix. *X_s_* contained the other significant jQTL peak markers from the interrogated chromosome and *X_S1_* is the genotype vector for each marker on the target chromosome within a window of 80 markers centered on the observed QTL peak being interrogated. LOD scores were computed as in Step 1 of “multitrait mapping to localize eQTL hotspots” and the position of the marker with the maximum LOD score was retained for each bootstrap resampling.

For all further analyses, we defined hotspot location confidence intervals as the central 95% confidence interval of bootstrap peaks. These intervals were extended to include all markers that are in perfect linkage disequilibrium with the markers at the ends of the confidence intervals.

### Local eQTL

To classify an eQTL as local, we required its location confidence interval to overlap the position of the gene. We used gene locations expanded by 1,000 bp upstream and 200 bp downstream to account for regulatory variants that may be located in the promoter or the 3’UTR. We initially classified 2,969 eQTL as local. These 2,969 eQTL affected 2,884 genes. Closer inspection revealed that these multiple “local” eQTL per gene often involved one eQTL with a peak very close to the gene and other, more distant eQTL. These more distant eQTL probably reflect trans eQTL on the same chromosome as their target gene with location confidence intervals broad enough that they happened to overlap the target gene. For our ASE comparisons below, we only used the local eQTL that were located closest to a given gene.

### Allele-specific expression analyses and comparison to local eQTL

We used ASE data from two sources. The first source is the mRNA data from Supplementary Data S2 from Albert et al., 2014 ^22^. The second source is unpublished data generated by Dr. Noorossadat Torabi in the Kruglyak laboratory. Both datasets performed mRNA sequencing on a BY/RM diploid hybrid strain. Reads from Albert et al. were ~30 bp in length to match ribosome profiling data presented in that paper, while reads from Torabi et al. were 100 bp. In contrast to ref ^22^, the Torabi data was not strand-specific.

We processed the Torabi data exactly as described in ref^22^. Briefly, reads were aligned to both the BY reference genome and the RM reference genome provided by the Broad Institute (https://www.broadinstitute.org/fungal-genome-initiative/saccharomyces-cerevisiae-rm11-1a-genome-project). We retained reads that mapped uniquely and without mismatch. We considered reads mapping to a set of coding SNPs carefully curated to only contain SNPs with good mapping characteristics^22^ and counted the number of reads arising from each allele. Counts for multiple SNPs per gene were summed. We performed hypergeometric downsampling to account for a small difference (0.5%) in total read counts mapping to BY vs. RM alleles. Genes with fewer than 20 reads were discarded. Significance of ASE was gauged using a binomial test, and p-values Bonferroni-corrected for multiple testing. Effect sizes are expressed as log2-transformed fold changes, which in turn are calculated as the RM allele count divided by the BY allele count for each gene.

The Torabi and Albert dataset are independent replicates of a BY/RM hybrid. We can use this fact to gain confidence in significance calls (we keep track of whether a gene was determined to have significant ASE in one or both of the datasets) and the magnitude of ASE (for each gene, we use the mean log2 fold change of the two datasets). Across the two datasets, we had ASE data available for 3,340 genes.

To compare effect sizes between eQTL and ASE, we used log2-transformed eQTL fold changes re-computed on un-scaled expression data. ASE fold changes were computed as the log2 of the ratio of RM to BY allele counts. Comparisons of effect size across genes were performed using standardized major axis (SMA) analysis^82^. To correlate ASE and eQTL effect sizes to the number of sequence variants upstream of each gene, we defined the upstream interval as the sequence upstream of the start codon up to the neighboring gene for a maximum of 1,000 bp. We acknowledge that the length of sequence considered is therefore different for different genes, but believe that this is a reasonable approximation of the regulatory upstream regions in yeast^83^. Sequence variants had been obtained from short-read sequencing of the BY and RM strains used in this study^15^.

All data necessary to reproduce the ASE analyses and eQTL comparisons is available in Supplementary Data File 7.

### Power analyses for ASE data

To gauge the statistical power to detect ASE in the two available ASE datasets, we performed simulations. We focused on two variables: the effect size (i.e. fold change) and the total read coverage available for the given gene. ASE data is overdispersed compared to a binomial distribution^84^. To properly account for this overdispersion in our simulations, we used the available ASE data to estimate the overdispersion parameter *ρ* in a beta-binomial distribution using the R function ‘optim’. We provided the function with the total read count and the observed allele count for each gene and used 0.5 as the “true” probability of success. We estimated p separately for the Torabi and the Albert data and found that the latter was somewhat more overdispersed (*ρ* = 0.0054) than the former (*ρ* = 0.0041).

We confirmed that these overdispersion estimates fit the data reasonably well by generating simulated datasets using the observed read counts from the Albert and Torabi datasets for each gene, along with the estimated ρ’s. For each gene, we used the mean Albert/Torabi ASE fold change to compute the target probability of success. These simulated data are meant to represent a random instance of allelic counts across genes, using distributions that closely mimic the real data. We generated 50 such datasets using the Albert and Torabi parameters, respectively. We computed all pairwise correlations between these simulated Albert/Torabi datasets. The median correlation coefficient across genes was *r* = 0.42 in the simulated data, compared to *r* = 0.39 for the observed Albert / Torabi data. Thus, while the simulations underestimate the overdispersion in the real data by a small amount, they provide a reasonable approximation.

We used these estimates for *ρ* to simulate 1,000 instances of allele counts for several combinations of total read count and fold change (Supplementary Figure 4). We computed power as the fraction of simulations in which a binomial test yielded p < 0.05 or p < 1.5e-5, the Bonferroni-corrected threshold for 3,340 tests. We also computed the fraction of simulations in which the direction of fold change agreed with the true change.

These simulations show that power increases with increasing read counts and effect sizes. Crucially, even with very high sequencing depth of 5,000 allele-specific reads per gene, the power to detect ASE of a magnitude typical for the majority of local eQTL detected here is less than 60%. Most genes in the ASE data have substantially lower coverages: the median coverage is ~1,000 in the Torabi data and ~200 in the Albert data. Thus, we expect to miss the ASE effects of the majority of local eQTL.

To explore this relationship more precisely, we conducted gene-matched power simulations. For each gene, we conducted 100 simulations using the estimated ρ and sequencing coverage for the given ASE dataset. We used the observed eQTL fold changes to compute expected probabilities of success. We conducted these simulations separately for the Albert and Torabi data.

### Genes affected by hotspots

To estimate which genes are influenced by a given hotspot irrespective of whether these associations reached genome-wide significance for the given gene, we performed a targeted forward scan for linkage at the hotspot locations. For each chromosome and for each gene, we regressed out the effects of significant QTL detected on other chromosomes as well as the effects of technical covariates. Then, for each gene, we repeated the procedure described above under “Mapping additive eQTL”. However, here *X* was defined to be the set of detected hotspot markers for that chromosome, as well as either the closest marker to each gene or, if a gene had a genome-wide significant local eQTL, the local eQTL peak marker for that gene. The same FDR threshold (5%) was used to identify the best model for each gene. For each gene, coefficients for the effects of all significant markers identified by this procedure were determined by multiple regression.

The regression coefficients from this model were used to count the number of genes affected by a given hotspot (i.e. all nonzero entries) and to rank genes for inclusion in gene ontology (GO) and transcription factor (TF) binding site (TFBS) enrichments, and for plotting in networks. Supplementary Data 8 also presents the number of genes with a genome-wide significant eQTL overlapping each hotspot.

For each hotspot, TFBS and GO enrichments were calculated on the 100 genes with the strongest increases or decreases in expression from the RM allele at the hotspot, respectively. When fewer than 100 genes were influenced by the hotspot in the given direction, we used all genes influenced in this direction. For the *CBF1* hotspot, the main text reports additional enrichment analyses of all 307 linked genes, which we conducted as a follow-up to the scan of the top 100 target genes.

In Supplementary Data 8, we report GO enrichments for the genes affected by each hotspot. The table shows the top five GO categories for each hotspot that exceeded an enrichment p-value of p ≤ 0.05 divided by the number of GO categories tested. We adjusted p-values separately for the Biological Process, Molecular Function, and Cellular Compartment GO subtrees. We did not adjust for the fact that multiple hotspots were tested, nor for the fact that two directions of effect (higher expression linked to the BY or the RM allele) were tested. Supplementary Data 12 lists all TFBS enrichment results.

TFBS enrichments for genes affected by hotspots were computed using Fisher’s exact test. The underlying relationships between TFs and target genes were based on regulatory relationships between genes downloaded from SGD^40^ on May 1^st^, 2016. In Supplementary Data 8, we show all TFs with significant enrichments at a threshold of p ≤ 1.3e-6, corresponding to a Bonferroni-corrected p-value controlling for 38,556 tests (189 TFs, 102 hotspots, and two directions of effect per hotspot). Supplementary Data 8 also shows the most significant TFBS enrichment, irrespective of whether this enrichment was significant after multiple testing or not. Supplementary Data 13 lists all TFBS enrichment results. For the *CBF1* hotspot, the main text reports additional TFBS enrichment analyses based on binding sites annotated by ref ^32^, using their “highest confidence sites”, defined as “those containing conserved motif matches that were bound by the corresponding factor at a p-value < 0.001”^32^, and downloaded from http://fraenkel.mit.edu/improved_map/. The resulting TFBS enrichment results for *CBF1* were very similar to those obtained using the annotation from SGD, which is a superset of the annotation in ref ^32^.

We displayed the genes that are affected by each hotspot in a manner that reflects significant correlations between the genes’ expression levels in order to highlight groups of genes that may be functionally related. Because expression levels will become correlated when they are linked to the same hotspot, we wanted to estimate the underlying correlation matrix in a manner that is as free as possible from correlations induced by the hotspots. We fit a sparse conditional Gaussian graphical model (sCGGM^85^) to the expression data and the genotypes at the hotspot markers. sCGGM decomposes the total correlation matrix among gene expression levels into “direct” effects of the genetic markers on each gene expression level as well as a matrix *θ_yy_* of “indirect” effects that genes exert on each other. We fit the model on a multi-core processor using an R installation compiled using the Intel Math Kernel Library to speed up linear algebra operations, as provided by the University of Minnesota Supercomputing Institute. Following preliminary cross-validation on a subset of 2,000 genes, we set the sCGGM regularization parameters *λ_1_* and *λ_2_* to 0.1.

We fit log2(TPM) values for all 5,720 expressed genes. Prior to fitting, effects of experimental batch and yeast culture optical density were removed using a linear model. We also removed the effect of the marker closest to each gene. This latter correction was not performed for genes that reside in the window tested by bootstrapping around each hotspot, and for genes with a local eQTL that overlaps the given hotspot. We chose to keep the local effects for these genes because local eQTL at these genes may underlie the hotspot, and we were interested in preserving such potential “direct” local effects for our visualizations.

We used the entries of *θ_yy_* to generate the network plots in Supplementary Figure 6 and Supplementary Data 11. In spite of the shrinkage imposed by the sCGGM algorithm, very few entries were estimated to be zero. As a practical threshold for plotting, we excluded values of *θ_yy_* with absolute values less than 1e-5.5. This threshold was set based on visual inspection of a histogram of all entries in *θ_yy_* which showed a bimodal distribution with a clear peak of values exceeding this threshold separated from a peak centered on much smaller values. Network plots were generated using the R igraph package^86^. Supplementary Data 11 shows the resulting network plots for all hotspots.

### Analysis of genes located in hotspots

Several hotspots were located close to chromosome ends. Yeast chromosome ends contain complex structural variation that segregates among isolates and influences traits ^87^. In some cases, BY and RM differ for the presence of entire subtelomeric blocks of genes ^88^. When a hotspot arises from these regions, the identity of the causal gene cannot be determined using our present segregant panel because each segregant either carries all or none of the genes in these regions. Further, the marker map we used for mapping stops at the borders of these regions. Therefore, these hotspots often have very sharp bootstrap distributions on the first or last marker of the linkage map on the given chromosome. We excluded subtelomeric hotspots from the analyses of genes located in hotspots because the position of the final marker on a chromosome is unlikely to reflect the position of the causal gene, which may well be located distally to the marker. We excluded 13 hotspots whose peak marker is within 5 kb of the end of our linkage map. We focused the remaining analyses of hotspot genes on 26 non-subtelomeric hotspots with confidence regions that contain three or fewer genes, for a total of 58 genes.

To analyze features of genes located in hotspots, we performed multiple logistic regression. The dependent variable indicates whether or not a gene resides in one of these hotspots, and the set of gene features described above served as potential predictor variables. Significance tests were performed using likelihood ratio tests for dropping each term from a full additive model without interactions, as implemented in the R car package.

Gene ontology analysis on these genes was conducted using Fisher’s exact test in topGO^74^. Genes in the yeast genome tend to be clustered in co-localized groups with similar functions^89^. To test if this clustering influences our GO analyses of genes located in hotspots, we performed a randomization analysis. We sampled 1,000 sets of 58 neighboring genes that mirror the distribution of the number of genes across the 26 hotspots. For each set, we performed the same GO enrichment analysis as for the actual 58 hotspot genes. Within each set, we counted the number of significant (at p < 0.05) GO terms and used the fraction of this distribution that matched or exceeded the number observed in the real set as an empirical p-value for whether the enrichment was globally significant. This test showed significantly more enriched terms than randomly expected for the GO Biological Process category (p = 0.001), but only a marginal excess for Molecular Function (p = 0.060) and no significant excess for Cellular Compartment (p = 0.695).

This analysis is conservative because in the real data we considered GO terms at much more stringent p-value cutoffs than ≤ 0.05. We further explored the FDR for GO term enrichment at more stringent p-values by dividing the mean number of terms significant at a given threshold across the permutations by the number of significant terms observed in real data. We found that for Biological Process, FDR was < 0.05 for GO enrichments with p < 0.005, which includes all terms described in the paper.

Finally, we computed an empirical p-value for each GO term by asking how often its observed p-value is matched or exceed in the permutations. This analyses controls for different sizes and compositions of the different GO terms. All terms reported to be significant in the text had p < 0. 001 in these analyses; enrichments as strong or exceeding the observed ones were never seen in 1,000 random gene sets. We conclude that our GO analysis of genes in hotspots is unlikely to reflect random sampling of genomic regions.

Plots of hotspot location and gene content were generated using the R package Gviz^90^. Supplementary Data 10 shows plots of gene content for all hotspots.

### Comparisons to pQTL

We used pQTL for 160 proteins identified in the BY/RM cross^18^. The pQTL coordinates were mapped from the sacCer2 to the sacCer3 genome using the UCSC liftover tool (https://genome.ucsc.edu/cgi-bin/hgLiftOver).

The pQTL were mapped using bulk-segregant analysis (BSA) in large pools of segregants where each gene was tagged with green fluorescent protein^18^. BSA does not produce effect sizes in units of gene expression levels or variance. We instead used the allele frequency difference between high and low GFP pools at the pQTL peak position as a measure of pQTL effect size. For eQTL, we used the coefficient of the correlation between scaled expression levels and marker genotype. We chose this effect size measure because it, like the BSA allele frequency estimates, is bounded by −1 and 1, resulting in more easily interpretable scatterplots. Using other measures of eQTL effect such as multiple regression coefficients did not change our conclusions about pQTL overlap.

For the comparison of strong eQTL to significant pQTL, we defined “strong” eQTL as those eQTL that explained ≥ 3.5% of phenotypic variance. Our eQTL data had ≥ 99% power to detect such eQTL. Under the assumption that the bulk-segregant based pQTL data had similar statistical power as the current eQTL data, these eQTL should be easily detectable as pQTL if their effects on proteins are similarly strong. For the comparison of strong pQTL to significant eQTL, we defined “strong” pQTL as pQTL with LOD ≥ 15. This threshold was chosen to pick a set of strong pQTL that had similar size (218 pQTL) to the set of strong eQTL (238 eQTL).

For eQTL that did not overlap a significant pQTL or vice versa, we used the effect size point estimate at the respective peak position in the non-significant dataset for plotting in Supplementary Figure 8 B & C and for results presented in Supplementary Tables 6 & 8. Data underlying these analyses is available as Supplementary Data 15.

### Genetic interactions

For each transcript with at least one significant additive QTL, we fit a model that included the batch and growth covariates, the significant additive QTL, and a random effect for polygenic background. The residuals from this model were used for the detection of QTL-QTL interactions.

The set of markers used for the detection of QTL-QTL interactions was reduced to 3,106 using the findCorrelation function in the R package caret (https://cran.r-project.org/web/packages/caret/index.html), using a cutoff of 0.99. All unique combinations of markers were tested for each transcript, with the exclusion of the 20 closest markers on the same chromosome. QTL-QTL peaks, including the rare case of closely linked QTL-QTL interactions occurring on the same chromosome pair, were identified using custom code. The same procedure was repeated for five random permutations of segregant identities. A false discovery rate was calculated as the ratio of expected to observed peaks at different LOD thresholds. A false discovery rate for the marginal scan was calculated as the ratio of expected to observed peaks at different LOD thresholds for the subset of marker pairs where one of the pairs had a significant additive effect. False discovery rate was controlled at 10% for both scans.

Additionally, we tested a model of QTL-QTL interactions between significant additive QTL only. For each gene, we regressed out the effects of significant additive QTL, the effect of polygenic background, and the effects of technical covariates. Then, for each gene we tested only the QTL-QTL interaction effect between significant additive QTL markers. We note that this procedure involved the peak markers detected in the section on eQTL mapping, and no marker downsampling was performed here. An F-statistic was calculated for each test. The same procedure was repeated 10 times with permutations of segregant identities. From the permutations, the expected number of significant QTL-QTL linkages was calculated at various thresholds. False discovery rate was controlled at 10% for this procedure. Supplementary Data 16 lists all identified genetic interactions.

## Acknowledgements

We thank Noorossadat Torabi for unpublished ASE data, Meru Sadhu for comments on the manuscript, and the Minnesota Supercomputing Institute (MSI) at the University of Minnesota for computational resources. This work was supported by funding from the Howard Hughes Medical Institute (to LK) and NIH grant R01GM102308 (to LK). FWA is supported by NIH grant 1R35GM124676-01.

## Author contributions

FWA, JSB, JS and LD performed the experiments. JSB and FWA analyzed the data. LK supervised the project. FWA, JSB and LK wrote the manuscript.

## Supplementary Figures

**Supplementary Figure 1.**
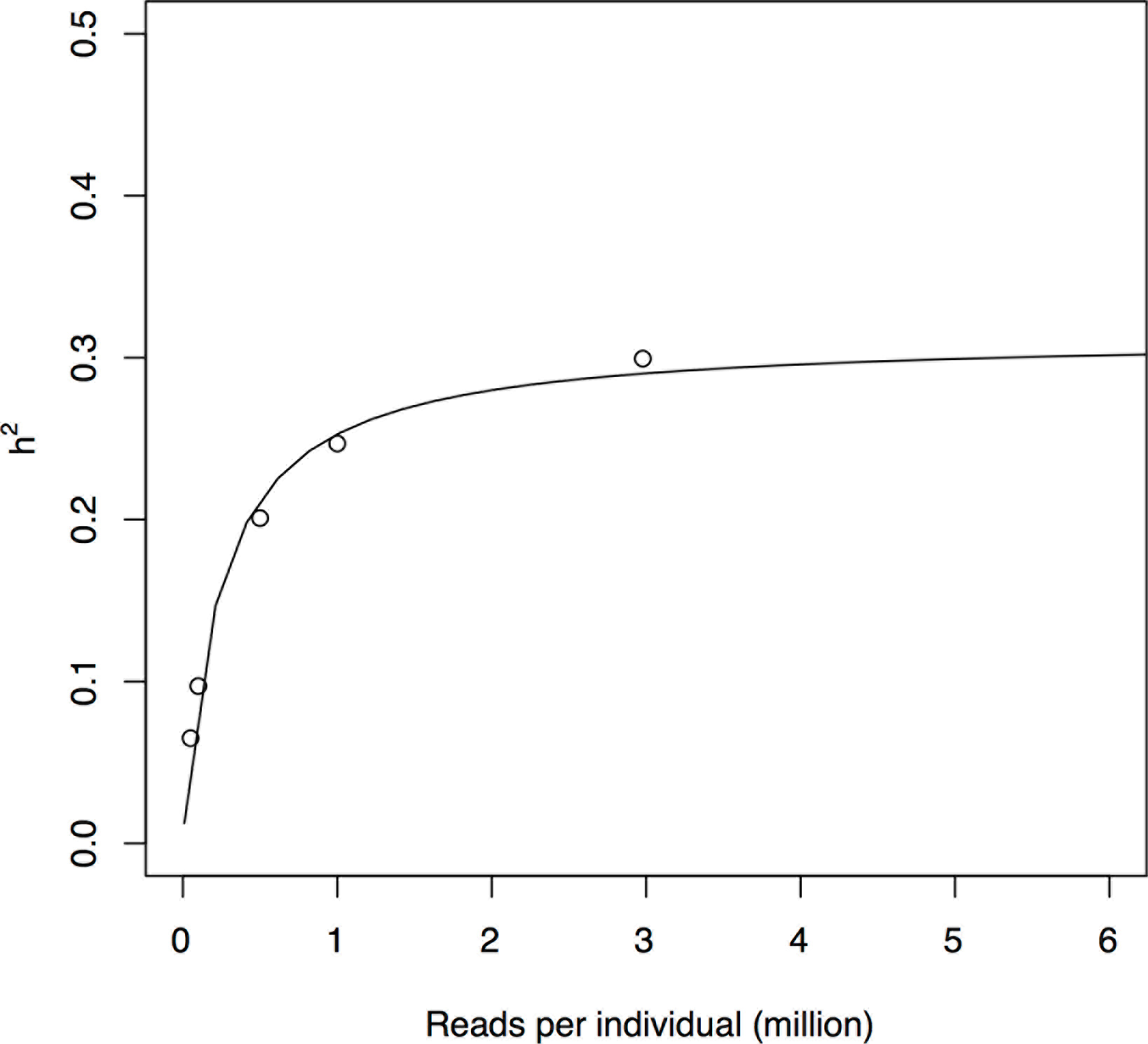
Mean additive heritability across all transcripts as a function of downsampling the total number of reads per sample. The downsampled number of reads per sample (x-axis) is plotted against the mean additive heritability across all genes (y-axis). The black line is a non-linear least squares fit.

**Supplementary Figure 2.**
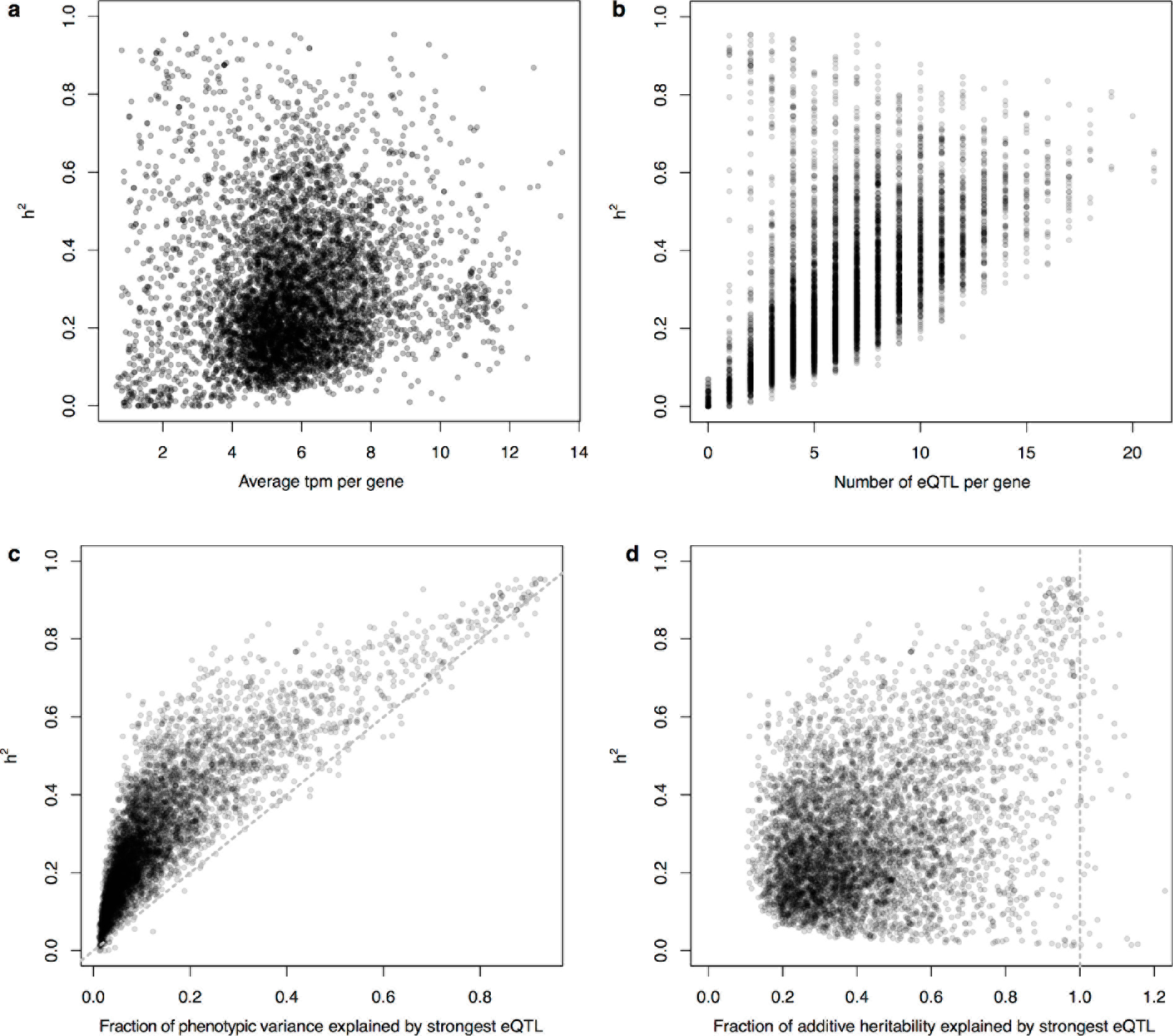
Heritability (h^2^) compared to other measures. In all panels, heritability for each gene is plotted on the y-axis and compared to: a) The average TPM for a gene across the segregants. b) The number of detected eQTL per gene. c) The fraction of phenotypic variance explained by the largest effect eQTL per gene. The diagonal line represents the case of all heritability mapping to the strongest eQTL, and is shown as a visual guide. d) The fraction of additive heritability explained by the largest effect eQTL per gene. The vertical line represents the case of all additive heritability being explained by the strongest eQTL, and is shown as a visual guide.

**Supplementary Figure 3.**
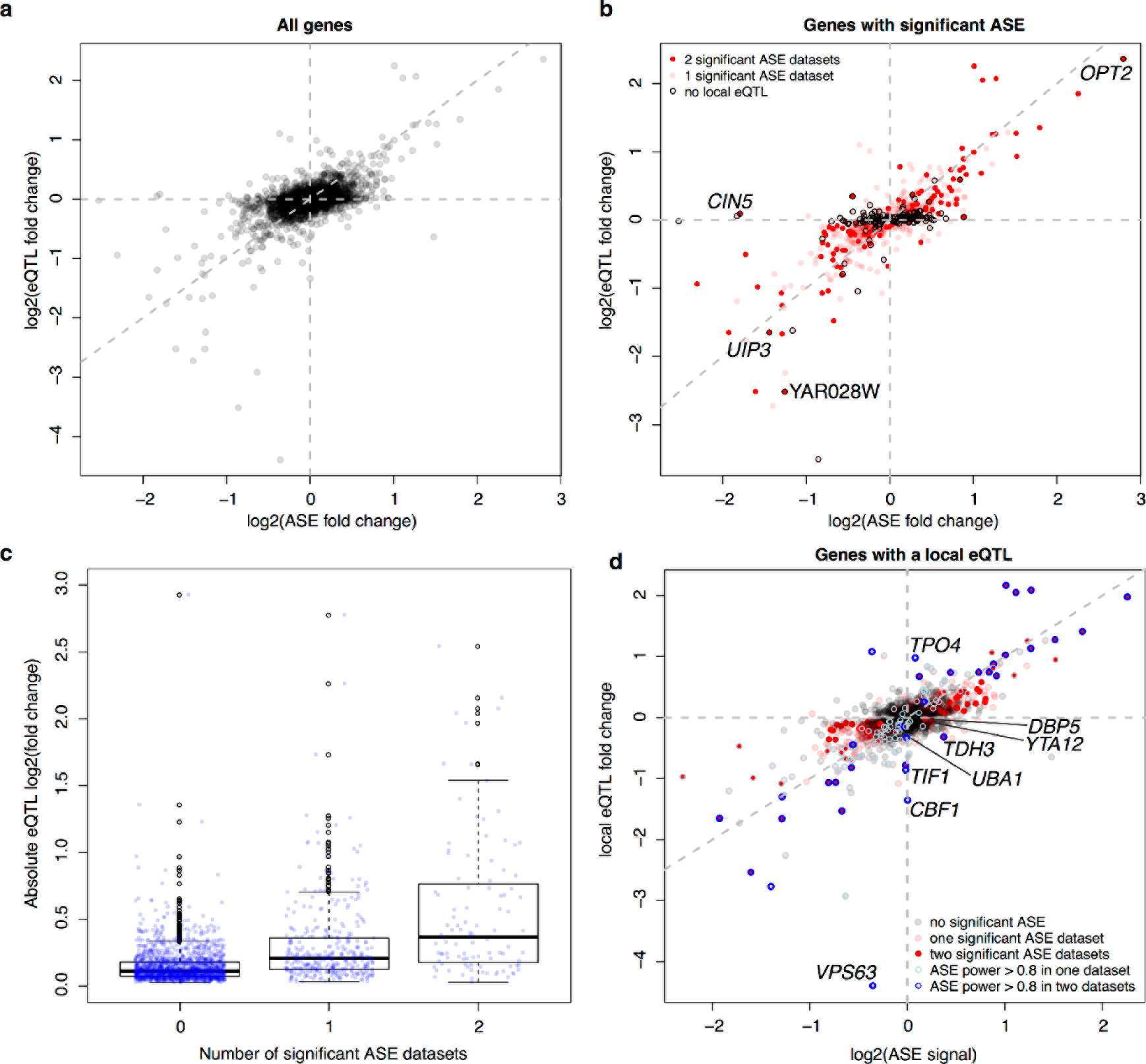
Allele-specific expression (ASE) compared to local eQTL effects. a) For each gene present in the ASE datasets, we show the magnitude of ASE in the diploid BY/RM hybrid (x-axis) vs. the magnitude of the local eQTL in the current data (y-axis). Positive values indicate higher expression in RM compared to BY. The vertical and horizontal lines indicate ASE and local eQTL effects of zero, respectively. The diagonal line represents identical ASE and local eQTL effects. Local eQTL effects are computed at the position of all genes, irrespective of whether the local eQTL was significant. b) As in a), but showing only genes with significant ASE in at least one ASE dataset. Genes without a significant local eQTL are highlighted by black circles. We show names of genes that have ASE in both datasets but do not have a significant local eQTL. c). Boxplots showing absolute local eQTL effects for genes with no, one, or two significant ASE datasets. d) as in a), but only for genes with a significant local eQTL. Genes with high statistical power to detect ASE are highlighted by blue circles. We show the names of genes with local eQTL and high ASE power but no significant ASE, and names of genes with significant ASE and a local eQTL with opposite direction of effect (*TDH3*, *YTA12*, *DBP5*).

**Supplementary Figure 4.**
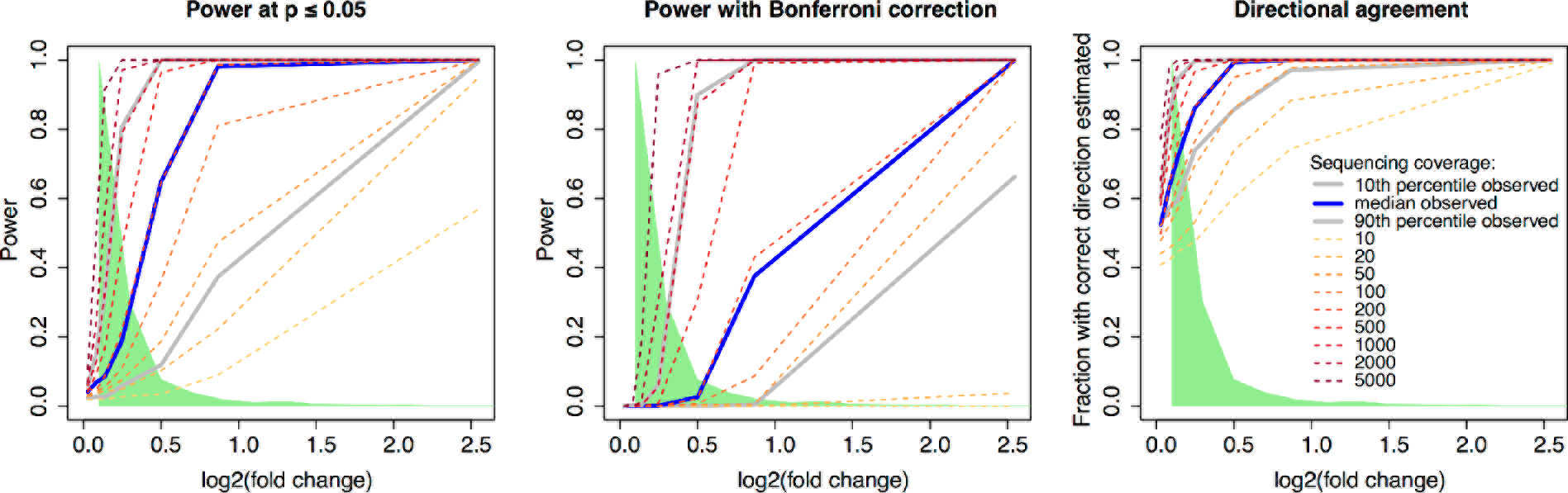
Power to detect Allele-specific expression. The figure shows the results from a simulation study that varied the strength of true ASE (x-axis) and the depth of sequencing coverage, expressed as the number of reads covering the two alleles of the gene. Different depths of coverage are shown as colored lines. The blue line indicates the median coverage per gene observed in ref ^22^, and the grey lines indicate the 10^th^ and 90^th^ coverage quantile in the same reference. The green area indicates the fold-changes observed for local eQTL to show the ASE magnitudes that may be expected in real data. On the y-axes, the panels show: Left: Power to detect ASE at nominal significance of p ≤ 0.05, Middle: Power to detect ASE with Bonferroni correction across the number of expressed genes, Right: the fraction of simulated ASE datasets in which the observed direction of ASE matched the true direction, irrespective of statistical significance.

**Supplementary Figure 5.**
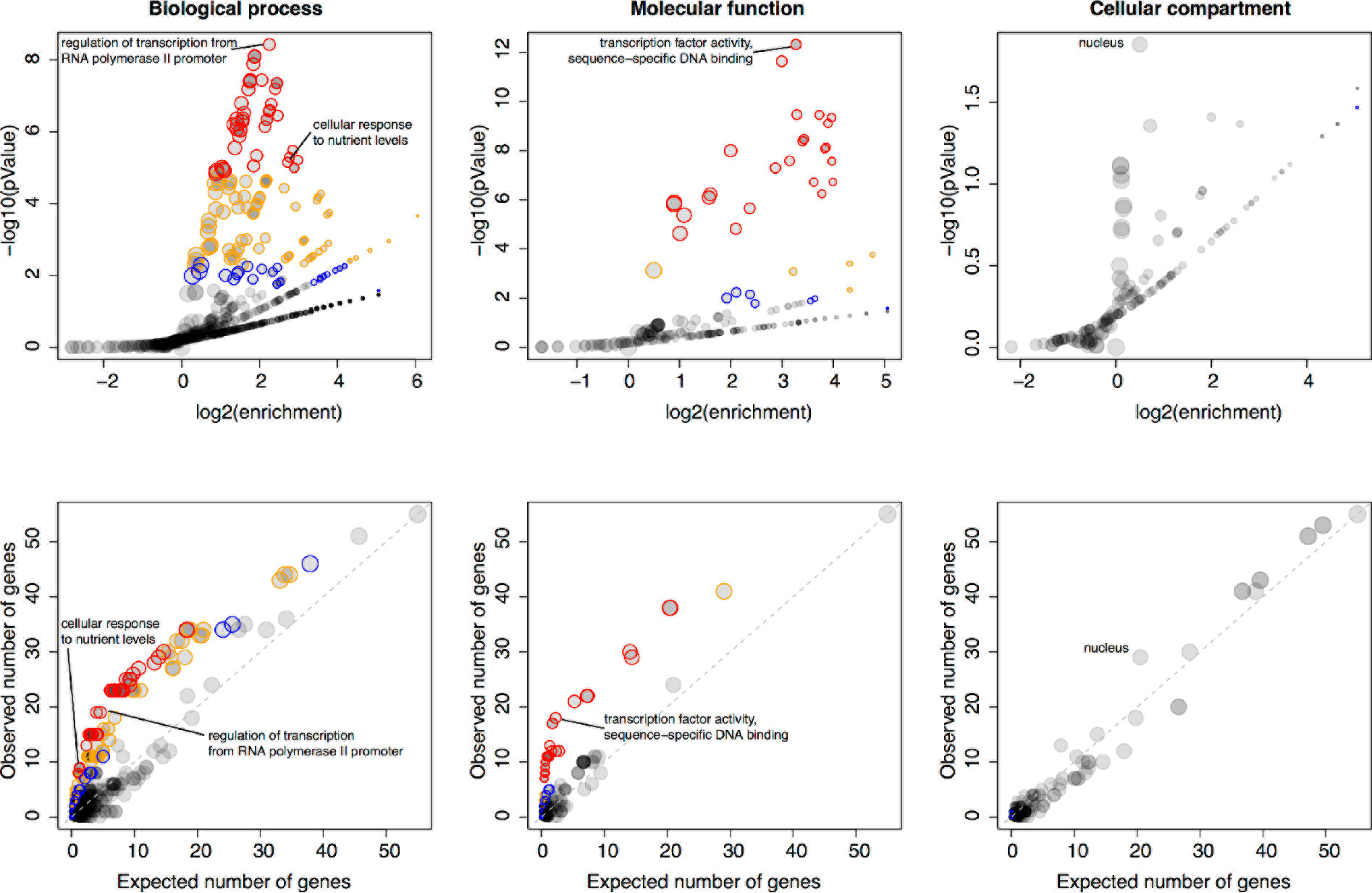
Gene ontology (GO) enrichments of genes located in hotspots. For each major GO category of “Biological Process”, “Molecular Function”, and “Cellular Compartment”, the figure shows two panels. The most significant GO term as well as GO terms discussed in the main text are indicated. Top panels: Relationship between strength (x-axis) and significance (y-axis) of the GO enrichment. Each GO term is plotted as a dot, with size scaled as a function of the number of terms in the GO group. Note how the relationship between enrichment strength and significance depends on GO category size. Different levels of significance are indicated by colored circles. With decreasing stringency, these colors indicate: Red: p < 0.05 after Bonferroni correction for the number of GO terms tested; Orange: permutation-based p < 0.005, corresponding to an FDR of 5% (Methods), Blue: GO term specific permutation-based p < 0.01. Bottom panels: The number of genes in each GO term expected to be significant based on GO category size (x-axis) vs. the number of genes in each GO term observed to be significant. Color codes are as in the top panels. The diagonal line indicates observations that match the expectation, and is shown as a visual aid.

**Supplementary Figure 6.**
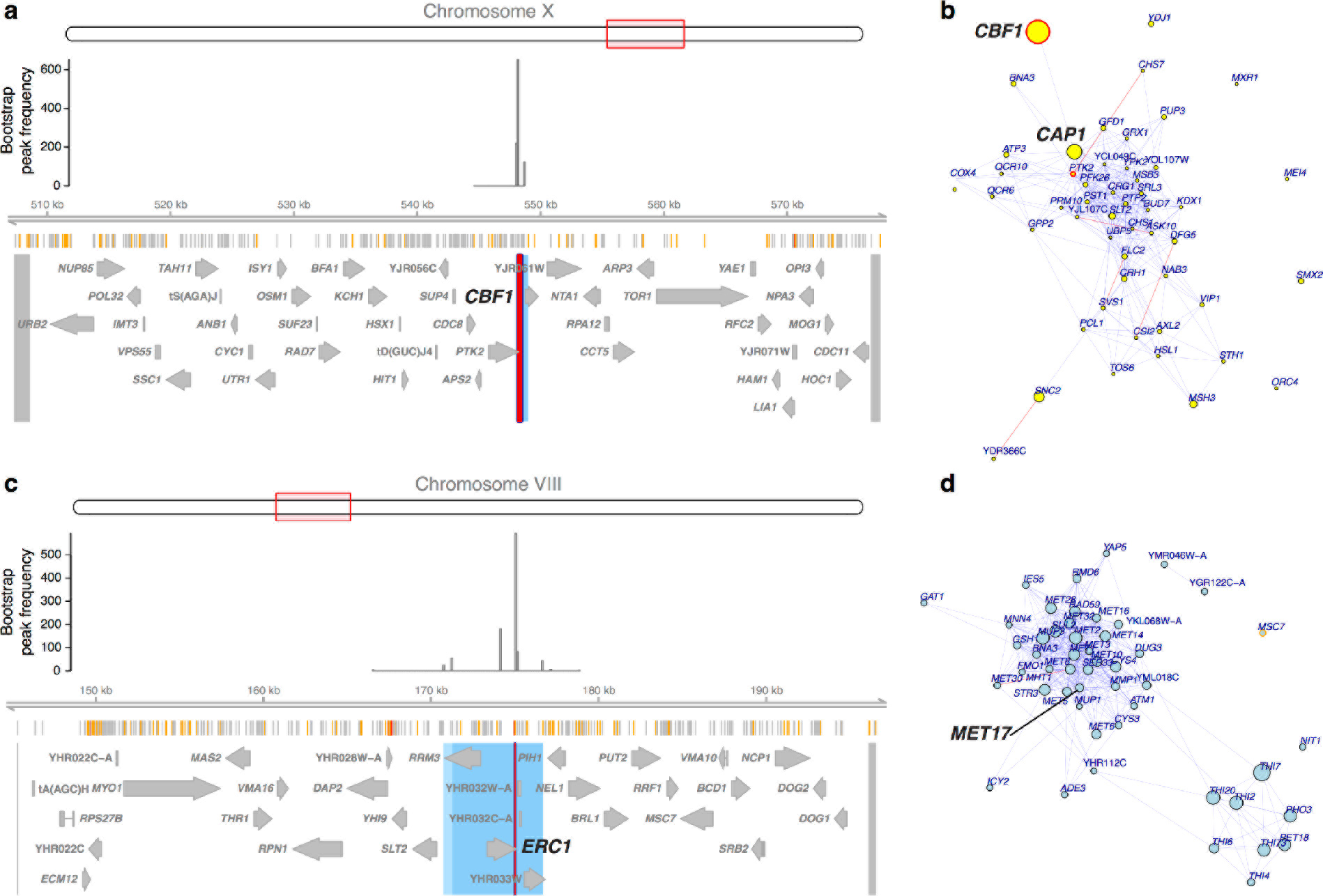
The *CBF1* and *ERC1* hotspots. a & c) The regions surrounding each hotspot. From top to bottom: a diagram showing the general region on the chromosome, the distribution of hotspot peak locations in 1,000 bootstrap runs, locations of BY/RM variants (red: variants with “high” impact such as premature stop codons^67^; orange: “moderate” impact such as nonsynonymous variants; grey: “low” impact such as synonymous or intergenic variants), and gene locations. The blue area shows the 90% confidence interval of hotspot location, and the lighter blue area shows the 95% confidence interval, both determined from the bootstraps. The red line shows the position of the most frequent bootstrap marker. The region tested in the bootstrap analysis is delimited by two markers, which are shown as grey lines at the outer edges of the plots. These markers and the peak markers are padded to span all variants that are in perfect linkage disequilibrium with the marker used in the analysis. b & d) A visual representation of the top 50 genes affected by each hotspot. Each gene is shown as a dot, with size scaled as a function of the size of the effect of the hotspot on the gene. For *CBF1*, we show genes with higher expression linked to the BY allele (yellow dots). For *ERC1*, we show genes with lower expression linked to the BY allele (light blue dots). Local eQTL that overlap the 95% confidence interval of hotspot location are indicated by red circles, and genes with a local eQTL anywhere in the region tested in the bootstrap analysis are indicated by orange circles. Edges between genes indicate co-expression in a gene regulatory network fit using code from ref ^85^. Blue edges indicate positive co-expression, and red edges indicate negative coexpression. Genes mentioned in the main text are highlighted. In c), note the connection between the local eQTL for *CBF1* and the distant eQTL for *CAP1*. In d), note the group of genes with methionine-related functions, including *MET17*.

**Supplementary Figure 7.**
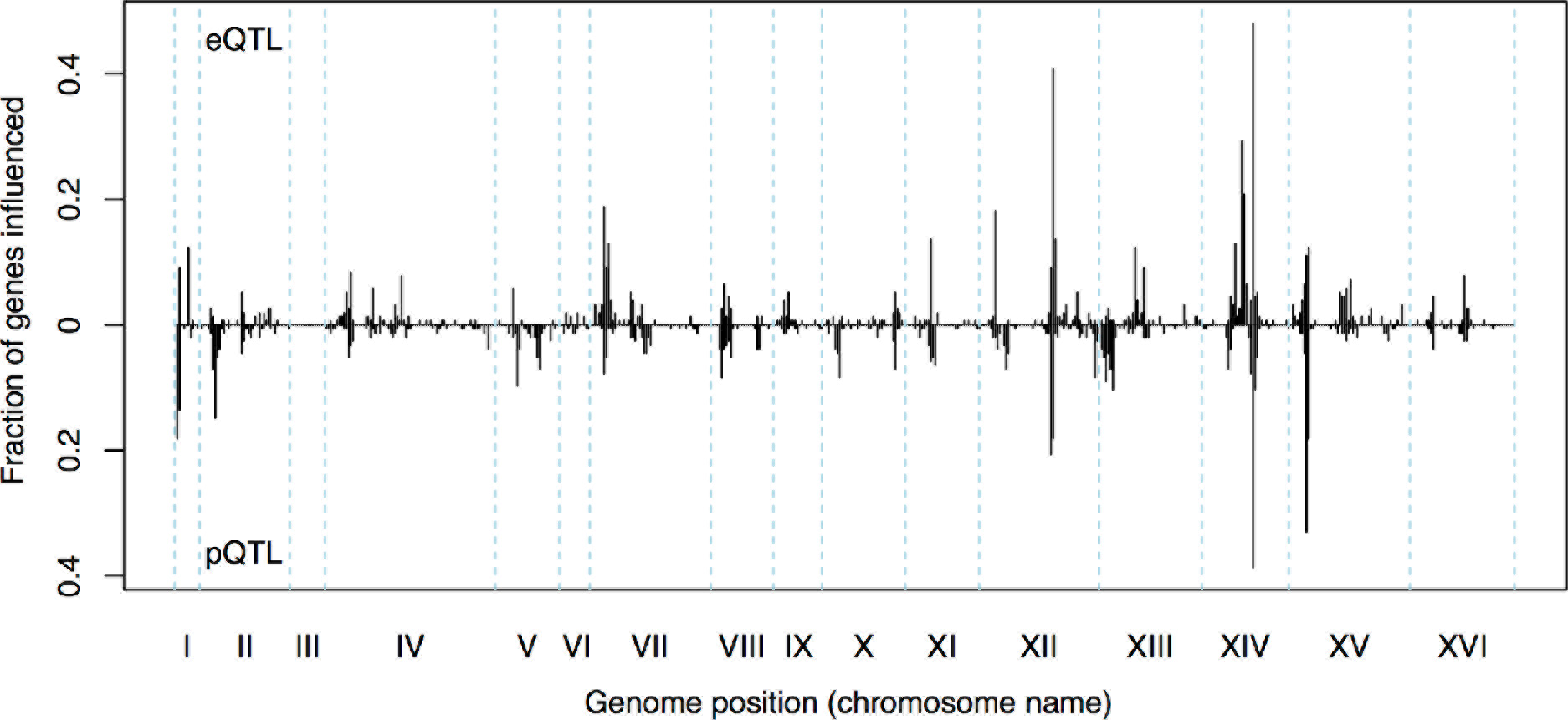
Distant eQTL and pQTL hotspots. The figure shows the fraction of 154 genes assayed for mRNA variation in the current data and for protein variation in ref ^18^ that have an eQTL or pQTL in a given bin along the genome (x-axis). eQTL from the current dataset are shown in the upper half of the figure, and pQTL from ref ^18^ are shown in the bottom half with an inverted scale. Note that chromosome III is omitted from the figure because eQTL hotspots on this chromosome cannot be detected as pQTL due the experimental design used to detect the pQTL^18^.

**Supplementary Figure 8.**
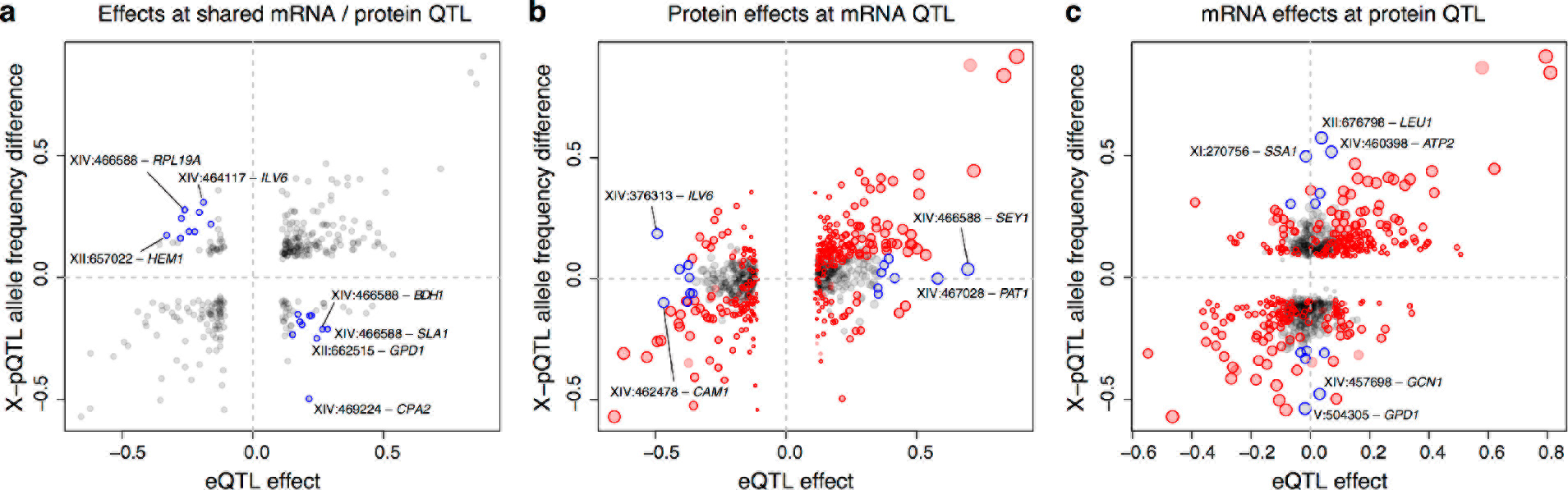
Comparison of individual distant eQTL and pQTL. Each panel shows the effect size of linkage of mRNA levels for a given gene to a given genomic position (x-axis; correlation coefficient between mRNA level and marker genotype) compared to the effect size of linkage of protein levels for the same gene to the same genomic position (y-axis; difference in frequency of the BY allele between pools with high and low expression of the protein as measured by green-fluorescent protein tags^18^). We chose these measures of effect size because they both can range from −1 to 1, facilitating visual comparison across different experimental designs. Positive values indicate higher expression in RM compared to BY. Only distant QTL located on different chromosomes than their target gene are shown. a) Effects at peak markers for overlapping significant eQTL and significant pQTL. Overlapping QTL with different direction of effect presented in Supplementary Table 7 are highlighted in blue; extreme outliers are marked in the figure by the genomic location of the QTL and the name of the affected gene. The dashed vertical and horizontal lines indicate zero eQTL and pQTL effect, respectively, and are shown as visual aids. b) At all distant eQTL, the figure shows mRNA and protein effects at the eQTL peak marker, irrespective of significance in pQTL data. Dot size scales as a function of eQTL effect size. Red circles denote eQTL that overlap a significant pQTL. Blue circles denote strong eQTL that do not overlap a pQTL as listed in Table Supplementary Table 6; the most extreme cases of eQTL without a pQTL are indicated. c) At all distant pQTL, we show mRNA and protein effects at the pQTL peak marker, irrespective of significance in eQTL data. Dot size scales as a function of pQTL effect size. Red circles denote pQTL that overlap a significant eQTL. Blue circles denote strong pQTL that do not overlap an eQTL as listed in Supplementary Table 8; the most extreme cases of pQTL without an eQTL are indicated.

